# Nanoscale precise stamping of biomolecule patterns using DNA origami

**DOI:** 10.1101/2025.05.09.653003

**Authors:** Laura Teodori, Ali Shahrokhtash, Elisabeth Asta Sørensen, Xialin Zhang, Mette Galsgaard Malle, Duncan Stewart Sutherland, Jørgen Kjems

**Author notes:** Correspondence should be addressed to Jørgen Kjems. These authors contributed equally to the work (Ali Shahrokhtash, Elisabeth Asta Sørensen).

## Abstract

Understanding the importance of ligand patterning in biological processes requires precise control over molecular positioning and spacing. While DNA origami structures offer nanoscale precision in biomolecule arrangement, their biological applications are limited by challenges related to their structural stability, scalability, and surface area. Here, we present a straightforward and rapid DNA origami stamping technique for transferring nanoscale oligonucleotide patterns onto surfaces, visualized using DNA-PAINT super-resolution microscopy to quantitatively assess the stamping efficiency and precision across different stamp types. Unlike traditional top-down methods that require specialized equipment, our technique provides an accessible, self-assembled platform for surface patterning, with versatility across various substrates via modifiable pattern-transfer oligonucleotides. We demonstrate reliable, efficient, and precise pattern transfer at single-molecule resolution, enabling new opportunities to study distance-dependent biological processes, including receptor activation, multivalent binding, and enzymatic cascades across broader spatial scales and different detection techniques. The use of the passivated surface limits non-specific interactions with unpatterned areas and enables control over the interaction between the biological target and the patterned biomolecules. Our method advances surface patterning by combining DNA nanotechnology with single-molecule imaging techniques, expanding access to cost-effective analytical approaches and potentially enabling multiplexed detection and live measurements.

---

Understanding how the density, geometry, and spacing of ligand patterns affect the response of receptors^1^ can be instrumental in studying and interacting with biological processes, such as cell migration,^2^ adhesion,^3^ and differentiation.^4^ These and many other biological phenomena can be investigated by placing biomolecules onto surfaces to recreate ligand patterns, a technique also known as biopatterning. Achieving this requires the precise positioning of biomolecules with geometries, densities, and intramolecular spacings that can be easily controlled and reproduced.^5^ Additionally, it should be capable of covering both small and large areas while being compatible with the biological system under investigation and with the technique used to detect changes in response to binding events.^5^

Several methods have been developed and are currently used to distribute biomolecules onto surfaces,^6^ with the most commonly used ones being direct-write patterning,^7–9^ lithography,^9^ and microcontact printing.^10,11^ These methods can offer high-throughput patterning of large surface areas with sub-100 nm resolution or better (the resolution of electron beam lithography can be below 10 nm).^12^ However, some patterning strategies may apply to only a few types of surfaces, limiting the number of applications and techniques capable of visualizing the pattern and/or detecting bioactivity. Finally, patterning methods may be time-consuming and not be globally accessible due to the need for expensive reagents and instrumentation.

The advent of DNA nanotechnology provided an alternative approach for surface patterning, as DNA origami structures can be implemented as nanoscale pattern elements or masks in lithography, or be functionalized and used as scaffolds for metal deposition, and mineralization. ^13^ Alternatively, two- and three-dimensional DNA structures have also served as a platforms to display ligands in a spatially- and stoichiometrically-controlled fashion, enabling studies of receptor-ligand interactions and their effects. DNA origami has been used to investigate means to enhance the affinity of receptor-ligand binding of cells by matching the avidity and spatial distribution of cell membrane receptors,^14,15^ observing changes in cell signaling according to ligand density,^16,17^ studying the viral receptor-binding domain distribution promoting cell infection and immune activation,^18–20^ and developing functionalized biosensors modulating density, orientation, and distance of target molecules to improve sensing performances.^21^ Despite these advances, the use of DNA origami as a platform to interface with biology can be easily compromised by its highly negative charge, which might interfere with the ability of the ligands to interact. Additionally, several key challenges remain unresolved, including the chemical and structural stability of DNA origami in biological settings and its limited surface area^22^ .

DNA origami can also serve as a molecular stamp to imprint patterns of molecules such as oligonucleotides and ligands onto surfaces.^23^ This approach has been demonstrated using both 2D and 3D DNA origami structures for functionalizing metal nanoparticles.^24–26^ Over the past decade, 2D origamis have been employed for large-scale surface patterning, particularly on gold substrates.^27–29^ In these applications, patterns of proteins or oligonucleotide displayed on the origami were transferred through either non-covalent (biotin) or covalent (-SH or -NH2 groups) interactions with a functionalized surface. The origami structure was subsequently removed from the surface through chemical cleavage or denaturation, revealing the stamped pattern. This approach is particularly promising as it leverages the ability to display precise nanoscale molecular patterns on DNA origami and replicate them across larger surfaces while eliminating the need for the structure in subsequent applications, thus avoiding its inherent limitations. While these stamped patterns were characterized by Surface Plasmon Resonance (SPR) and visualized by Atomic Force Microscopy (AFM), quantitative analysis of the efficiency and precision of the stamping remains limited. Moreover, the restriction to gold surfaces has constrained the use of other powerful techniques such as single-molecule localization microscopy (SMLM), which has become instrumental in the study of biological phenomena.

In this study, we demonstrate a novel approach using DNA origami as a stamp to transfer nanoscale patterns of oligonucleotides onto a polyethylene glycol (PEG)-coated glass-based coverslip, allowing the implementation of the super-resolution SMLM technique, DNA point accumulation for imaging in nanoscale topography (DNA-PAINT).^30^ This approach not only allowed us to visualize individual stamped oligonucleotides but also enabled a quantitative assessment of patterning efficiency and precision across different immobilization strategies, highlighting their respective strengths and limitations. Moreover, the dense PEG brush confers stealth to the coated glass surface, minimizing non-specific binding and protein adsorption, thereby facilitating future studies on ligand-receptor binding, including cell-based interactions, while also enabling applications that require controlled molecular geometries and distances.

## Results and Discussion

### Design of the stamping mechanism

The DNA origami stamp was designed as a double-layered rectangle to confer stiffness to the flat surface of the structure and minimize distortions of the geometry of the pattern during the stamping process (Figure 1 A).

**Figure 1.**
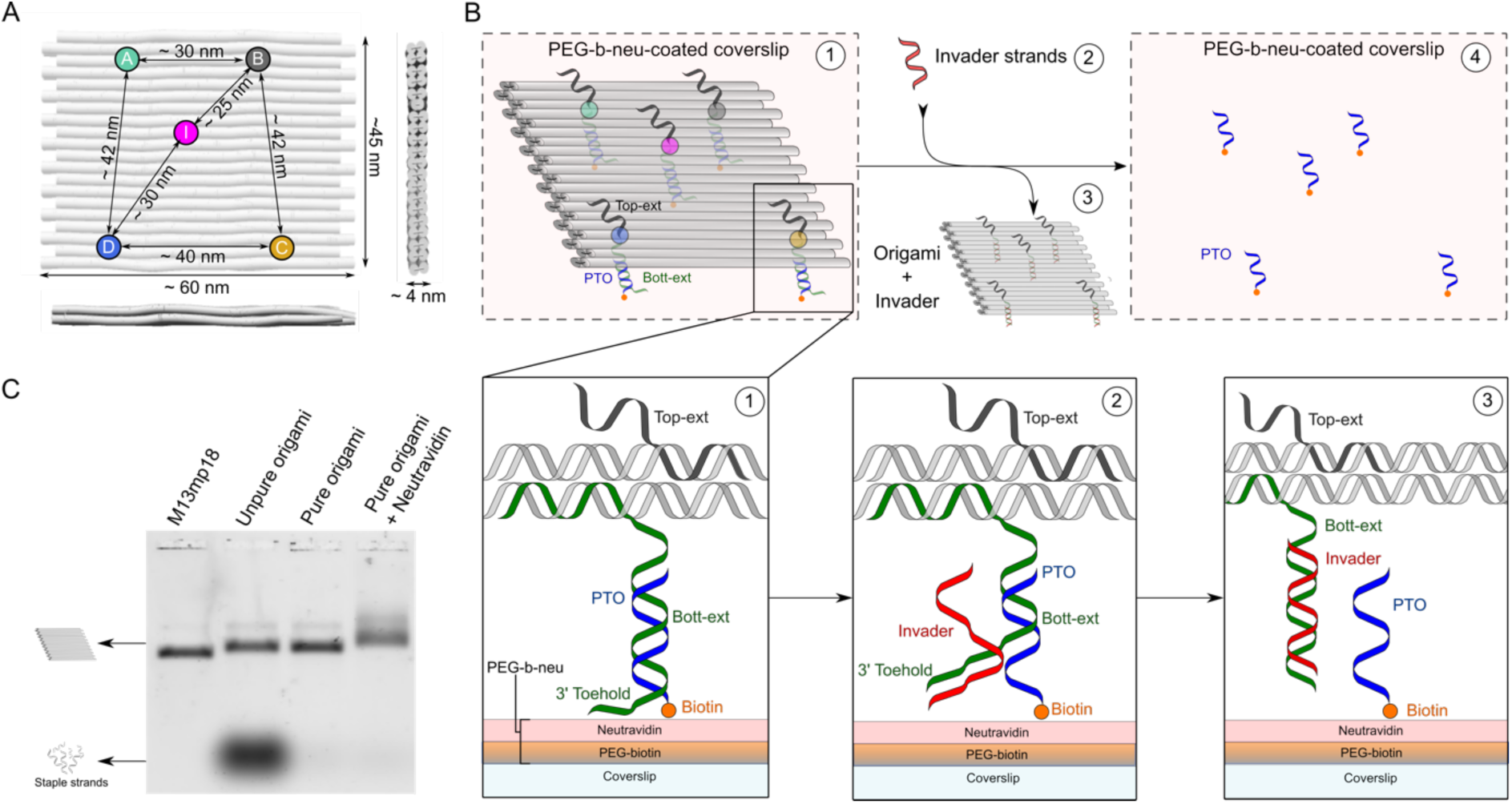
Molecular stamping mechanism design. (A) CanDo^31,32^ rendering of the DNA origami stamp edited to represent the dimension of the structure and the distances between the spots constituting the vertices of the trapezoid to be stamped. (B) Stamping mechanism design: 1) The origami stamp is immobilized on a PEG-b-neu surface through the biotinylated PTOs annealed to the staple oligonucleotide extending from the bottom of the DNA structure, denoted Bott-ext. The Top-ext staple oligonucleotide is used to visualize the origami stamp immobilized on the surface (described in the next section) 2) The invader oligonucleotide is introduced to trigger the toehold-mediated PTO displacement, and 3) the origami stamps are released in solution, which is then washed off. 4) The stamped PTOs remain immobilized on the surface through the established biotin-neutravidin interaction as a result of the pattern stamping process. (C) Agarose gel electrophoresis of the M13mp18 scaffold and the assembled origami stamps before and after purification. The purified structures were successively incubated with excess neutravidin and run in the gel. The upward band shift confirms that the biotin-PTOs are annealed to the origami stamps and available for neutravidin capture.

The origami stamp measured approximately 60 nm x 45 nm and displayed a trapezoidal pattern of staple strand extensions protruding from the top and bottom surfaces. The trapezoid’s vertices and an internal point were denoted respectively as A, B, C, D, and I, with inter-distances ranging between 25 and 42 nm. From each point constituting the pattern, the two staple strand overhangs were extended in opposite directions (Figure 1 B-1): one protruding from the top (denoted as Top-ext) and the other from the bottom surface of the structure (denoted as Bott-ext). The Top-ext remained unpaired, as it was intended for the detection of the origami stamp immobilized on the surface in the first step of the stamping mechanism. The stamp and the resulting stamped patterns were validated by DNA-PAINT technology, providing super-resolved visualization of the single oligonucleotides and enabling qualitative and quantitative characterization of the stamping process. The Bott-ext was annealed to a pattern transfer oligonucleotide (PTO), which served a dual purpose: immobilizing the origami on a surface and acting as the ‘ink’ released during stamping. The 5’ end of the PTO can be functionalized with various anchoring groups, such as biotin or a reactive group (-COOH, -SH,-N_3_, etc.) that bind the surface non-covalently or covalently. Additionally, the design of the Bott-ext-PTO complex enables the interchange of functionalized PTOs on the same assembled stamp without significantly altering the core structure of the origami. This feature allows the PTO to be annealed to the structure either during or after the assembly of the origami stamp, adapting the functional group and thus stamping mechanism to virtually any relevant surface, and making the testing of multiple conditions more time- and labor-efficient. In this study, the 5’ end of the PTO was functionalized with biotin to allow the immobilization of the origami structure and the stamping of single PTOs onto a dense neutravidin-coated surface. The functionalized surface denoted as PEG-b-neu (Figure 1 B-1) was prepared on a coverslip to enable fluorescence microscopy imaging. It consists of a monolayer brush of poly(acryl-amide)-g-PEG-biotin polymer ^33–35^ as ‘stealth’ low non-specific binding background, successively coated with a high concentration of neutravidin to maximize the density of available binding sites.

The stamping mechanism requires that the origami stamp is stably captured on the neutravidin-coated surface by the biotinylated PTOs annealed to the Bott-ext of the stamps (Figure 1 B-1). The transfer of bound PTOs from the origami onto the surface is triggered by a toehold-mediated strand displacement mechanism^36^. Here, an invader oligonucleotide is introduced into the system (Figure 1 B-2), which anneals to the toehold sequence located at the 3′-end of the Bott-ext. This leaves each PTO attached to the surface while releasing the origami in solution (Figure 1 B-3). Subsequent washing of the surface removes the origami stamps and excess invaders, revealing the stamped nanoscale pattern of PTOs stably anchored to the PEG-b-neu surface (Figure 1 B-4).

The core origami stamp was assembled, and its structural quality was assessed by transmission electron microscopy (TEM) (Figure S1). Once the biotinylated PTOs were annealed to the core structure, the integrity of the stamp was assessed by agarose gel electrophoresis (Figure 1 C). The presence of biotinylated PTOs was confirmed by the upward shift caused by the binding of neutravidin added in excess to the purified PTO-functionalized origami.

### Characterization of the molecular stamping mechanism

To verify the functionality of the PTO displacement mechanism, the ability of the invader oligonucleotide to trigger toehold-mediated strand displacement was tested (Figure S2). Each of the five Bott-ext oligonucleotides alone was annealed with biotinylated PTOs to form Bott-ext-PTO complexes. Subsequently, excess invader oligonucleotide was added to the complexes to trigger strand displacement, release PTOs, and form Bott-ext-invader complexes. The successful PTO displacement was visualized by native gel electrophoresis and confirmed by the slower migration of each of the Bott-exts-invader complexes formed upon strand displacement, compared to the migration of single Bott-ext oligonucleotides alone or those annealed to the PTOs.

An initial characterization of the origami-mediated molecular stamping mechanism on the PEG-b-neu surface using assembled origami stamps was performed by SPR (Figure S3). The increased response caused by the origami immobilizing on the surface, rapidly decreased upon the addition of the invader sequence, which released the origami from the surface. Upon washing the surface, the persistence of a residual response indicated the presence of stamped PTOs, which were still bound to the surface.

To characterize the molecular stamping mechanism visually, DNA-PAINT^30^ docking sequences 3xR3 and 3xR1^37^ (3xR3d and 3xR1d) were respectively incorporated within the sequences of the Top-exts and PTOs to enable super-resolved DNA PAINT imaging of the origami stamps and the stamped patterns by total internal reflection fluorescence microscopy (TIRFM). Multiplexed imaging of the two targets on the same field of view (FOV) was performed by the exchange-PAINT technique,^38^ where the two corresponding dye-labeled imager strands are sequentially introduced to transiently bind to their target docking strand.

In the first imaging step (Figure 2 A-1), the R3 imager strand conjugated to the Cy3B fluorophore (R3i-Cy3B) and complementary to the concatenated sequences of 3xR3d in the Top-exts, was applied to visualize the origami immobilized on the PEG-b-neu coverslip surface.

**Figure 2.**
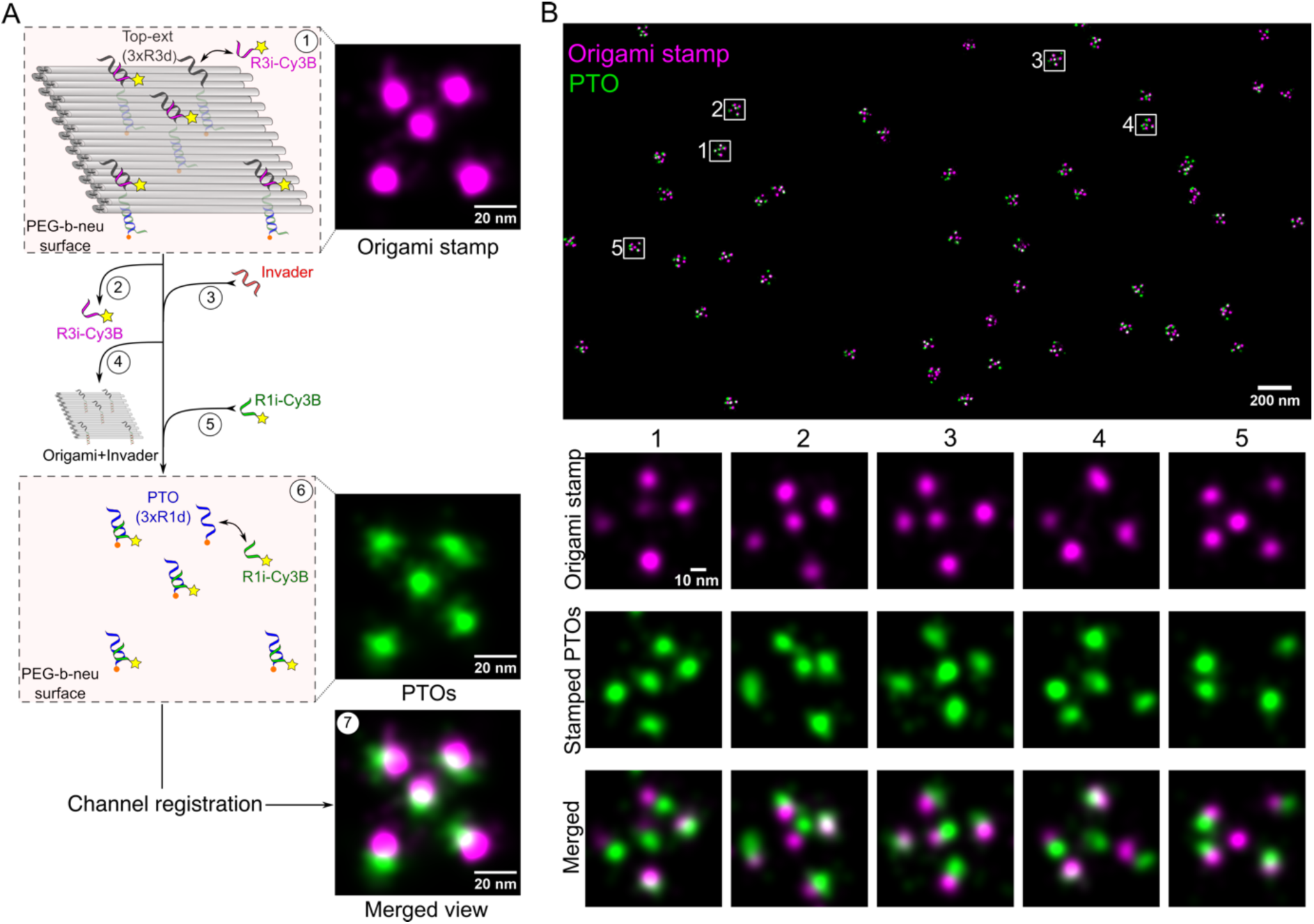
DNA-PAINT visualization of the stamping mechanism. (A) Schematics of the imaging steps in the stamping process. The immobilized origami stamp situated on the PEG-b-neu-coated surface was visualized by DNA-PAINT through the application of the R3i-Cy3B imager, which transiently anneals to the Top-exts of the origami (1). After the first imaging, the R3i-Cy3B imager is washed off (2), and an excess of invader oligonucleotides is applied to trigger the strand displacement of the PTOs (3), releasing the origami stamps in solution (4). The R1i-Cy3B imager, complementary to the 3xR1d sequence on the PTOs, is applied (5) to visualize the single-stranded PTO sequences immobilized on the surface upon the stamping mechanism (6). Upon image processing and channel registration, the origami stamp and PTO images are merged to locate each printed pattern. (B) In the top panel, a magnified ROI visualizes the signals merged by Picasso Render software of the origami stamps and the corresponding stamped PTOs. In the lower panel, the zoomed-in regions, including five origami stamps and their relative stamped PTO patterns, are shown as individual and merged signals. The local alignment of the picks was performed by subtracting the centroid for each channel, thereby centering both channels around the weight of the individual picks.

Following the origami stamp visualization, the R3i-Cy3B imager was washed away (Figure 2 A-2), and an excess of invader oligonucleotides was applied and incubated onto the surface for strand displacement (Figure 2 A-3). The subsequent washing step removed the excess invader sequence and the origami stamps in solution (Figure 2 A-4). The addition of a second imager strand, R1i-Cy3B (Figure 2 A-5), enabled its transient binding to its corresponding docking sequence in the PTO. Notably, the 3xR1d sequence incorporated in the PTO becomes single-stranded and accessible for imager docking only after strand displacement (Figure 2 A-6). This ensures that only single-stranded PTOs immobilized on the surface are visualized, revealing the intended stamped pattern. The signals from the origami stamps and single-stranded PTOs, acquired within the same FOV, were overlaid using gold nanoparticles added to the surface as fiducial markers. The resulting view (Figure 2 A-7 and S4 A, B, and D) revealed that the visualized stamped PTO pattern corresponded to each detected origami stamp. Signals generated by the origami stamps and their corresponding stamped PTO patterns are easily distinguishable from pattern-unrelated signals. Therefore, their localizations were visually identified and filtered to obtain the view in Figure 2B (top panel). The sparse single spots, highlighted by the white arrows in Figure S4 B and D, and removed in the filtering process, suggested that the PTO signal might have originated from free PTOs, that were not annealed to the origami and were not successfully removed during origami purification. The successful removal of the origami stamps was assessed by reapplying Exchange PAINT. The Top-exts of the origami stamps were visualized after strand displacement by reapplying R3i-Cy3B. The resulting images in Figure S4 C, F, and G confirmed that only a small number of origami stamps remained attached to the surface after incubation with the invader oligonucleotide.

The high average localization precision measured by Nearest Neighbor Analysis^39^ (NeNA, Table S1) achieved by DNA-PAINT imaging (Table S1) allowed to successfully resolve distances among the single points of the pattern displayed by each origami stamp and the stamped PTOs (Figure 2 B, bottom panel). Additionally, this technique enables a controlled real-time monitoring of the stamping mechanism in a fixed FOV directly on the microscope stage, potentially facilitating further experiments involving DNA-PAINT or other SMLM techniques and enabling visualization of how the newly formed pattern affects the biological phenomenon under analysis.

By visually inspecting individual patterns, we observed that while the overall pattern of the origami stamp closely matched the designed reference, the distances between the stamped PTOs varied among the individual stamps. To characterize these differences, alternative immobilization strategies were tested to assess changes and identify potential improvements for enhancing stamping efficiency (defined as the number of stamped PTOs for each origami), precision, and accuracy in reproducing the geometry of the pattern.

### Stamping efficiency of mono- and trivalent Biotin-PTO-based stamps

Efficiency and precision are important parameters for a surface patterning technique. However, the efficiency of the molecular stamping system is highly dependent on the surface to be patterned, the PTO properties, as well as on the features characterizing the origami stamps, such as structural rigidity. The stamping mechanism was tested on the PEG-b-neu surface to investigate and characterize its efficiency when the PTO was coupled to either one or three biotin molecules (Figure S5 A and B), respectively generating Biotin-PTO and 3xBiotin-PTO (Figure 3 A). Furthermore, a longer PTO, with a 7-nucleotides 5′ T spacer (7T-PTO, Figure 3 A), was also conjugated to three biotins (3xBiotin-7T-PTO, Figure S5 A) to maximize the flexibility and chances of PTOs of binding to neutravidin.

**Figure 3.**
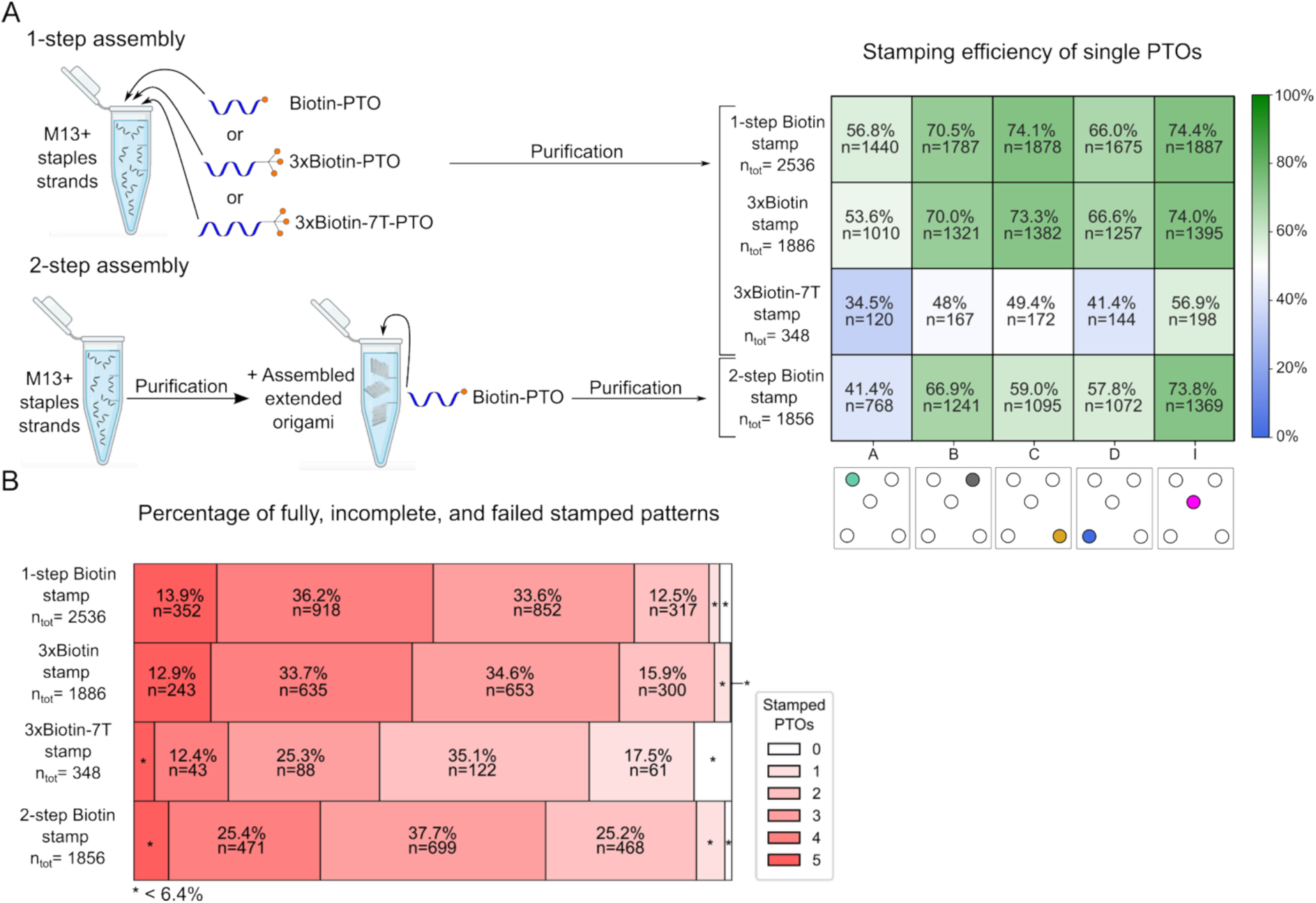
Stamping efficiency. (A) The origami stamps were assembled through either a 1-step or 2-step assembly. In the 1-step assembly, the M13 scaffold, the core and extended staple strands of the origami, and either biotin-PTO, the 3x Biotin PTO, or the 3x Biotin-7T PTO, were mixed, assembled into the relative origami stamp, and purified. In the 2-step assembly, only the M13 scaffold, along with the core and extended staples, were mixed and assembled to the origami structure. After a first purification, the monovalent biotin-PTO was annealed to the Bott-exts of the assembled extended origami structure, followed by a second purification to remove excess PTOs. The stamping process of each was evaluated by DNA-PAINT imaging and their stamping efficiency was reported as either the frequency of stamping of each PTO (A, B, C, D, and I) or (B) as the percentage of fully (5 stamped PTOs), incomplete (with 4,3,2, or 1 stamped PTOs), or failed (0 stamped PTOs) stamped pattern. All the percentages and the number of observed patterns for each origami stamp type are reported in Table S2. The total number of analyzed origami stamps and corresponding stamped patterns is indicated by “n_tot_”, whereas “n” indicates the number of observed stamped PTO (for right panel in A) or stamped patterns (for panel B) relative to the reported percentage. The results are based on one representative experiment for each type of origami stamp.

The origami stamps were assembled via either a 1-step or 2-step protocol (Figure 3 A). In a 1-step assembly, the origami stamp was assembled by mixing the M13mp18 (M13) scaffold, the core staples, the Top-ext and Bott-ext staples, and either Biotin-PTOs, 3xBiotin-PTOs, or 3xBiotin-7T-PTOs, followed by a single purification to yield the final origami stamp. This approach was found to maximize the number of annealed PTOs, thereby improving the overall stamping efficiency. In the 2-step assembly, the origami stamp was first assembled without any PTOs by mixing the M13 scaffold, the core, and the extended staples. The origami was then purified, incubated to anneal with excess Biotin-PTOs, and re-purified to obtain the final origami stamp. This approach enables modularity by first constructing the core structure and, at a later stage, adapting it to the specific surface by adding the relevant PTOs. Additionally, this approach reduces the excess amount of PTOs needed for the assembly and the residual non-annealed PTOs in the solution.

To characterize and quantify the efficiency of the molecular stamping performed by the 1-step Biotin stamps, 3xBiotin stamps, 3xBiotin-7T stamps, and 2-step Biotin stamps, DNA-PAINT localization events belonging to the Top-exts of the origami stamp and the relative stamped PTO have been aligned onto a simulated pattern model and clustered (Figure S7 B). Each cluster was then classified as position A, B, C, D, or I (Figure S7 C), and the stamping efficiency of the single PTO positions (Figure 3 A, right panel) was evaluated, along with quantification of the number of fully (5 PTOs), incomplete (4, 3, 2, or 1 PTOs), and failed (0 PTOs) stamped pattern (Figure 3 B). A detailed description of the data processing and analysis workflow is elaborated in the Materials and Methods section.

The data indicated that the stamping efficiency depended on the PTO position. Position I recorded the highest frequency of stamping across the four different types of origami stamps used, followed by the PTOs at positions C, B, D, and A (Figure 3 A, right panel). This may suggest that the surface of the origami stamp might not be completely flat when in solution. Instead, the central and right portions of the origami, including PTOs at positions B, C, and I, could be slightly concave. As a result, PTOs displayed at these positions could protrude more prominently than PTOs at positions A and D do. This may increase their proximity to the surface and the chances of binding to neutravidin.

The 1-step biotin stamp showed the best transfer efficiency for four out of the five PTO positions (Figure 3 A). Furthermore, its overall transfer efficiency was the highest for fully transferred patterns (13.9%, Figure 3 B), with a higher frequency (respectively, 36.2% and 33.6%) of stamped patterns displaying 4- and 3-PTOs. The use of 2-step Biotin stamps substantially reduced the frequency of fully transferred patterns down to 5.8% (Table S2) while predominantly increasing the stamping of 4-, 3-, and 2-PTO patterns. This demonstrated that the 1-step assembly protocol, where a temperature gradient is used, greatly increased the number of PTOs annealed to the origami stamp. However, the 2-step assembly could be a convenient approach for achieving acceptable surface patterning, enabling fast screening of multiple surfaces using the same stamp and smaller amounts of various functionalized PTOs.

Compared to the 1-step biotin stamp, the stamping efficiency did not substantially improve when three biotins were conjugated to each PTO (Figure 3 B and Suppl. Figure S6 A, B, C, and D). While the frequency of fully and incomplete stamped patterns was only similar to those observed for the 1-step biotin (12.9%), the percentage of failed stamped patterns was reduced to 0.2% (Table S2), indicating that the 3xBiotin stamp consistently guarantees the stamping of at least one PTO. This suggests that while the conjugation of three biotin molecules increased the likelihood of successfully immobilizing more PTOs, it was not sufficient to enhance the overall number of fully stamped patterns.

Coupling three biotin molecules to a longer and more flexible 5′ end of the PTO did not improve the stamping efficiency. Instead, it negatively impacted the frequency of fully stamped patterns by reducing it to 3.4% and increasing failed stamping to 6.3% (Table S2). Given the retained efficiency of strand displacement in removing the majority of the origami stamps (Figure S6 E, F, G, and H), this result may be ascribed to the 7T-linker extension (∼2.3 nm), which could have reduced the number of 3xBiotin-7T-PTO strands annealed to the Bott-ext or limited the ability of 3xBiotin-7T-PTO to bind to the neutravidin surface when complexed with the origami stamp.

### Stamping precision and accuracy of the origami stamps

Improving the efficiency of stamping enables testing of as many complete stamped patterns as possible, while simultaneously optimizing the stamping precision ensures that the potential ligands are precisely and consistently spaced at the intended distances, faithfully reproducing the desired pattern on the surface.

A qualitative characterization of the precision of the stamping system in reproducing the designed pattern is displayed in Figure 4A, where all the observed patterns from the origami stamp through the Top-exts and their relative stamped PTOs are represented as heatmaps. The Pearson correlation coefficient was used as a parameter to estimate stamping precision, and it was calculated between the DNA-PAINT data collected for each origami stamp and its corresponding stamped patterns. The Pearson correlation values revealed that the 3xBiotin origami offered a stronger stamping precision compared to the other structures (Figure 4A). The correlation measured for the 1-step and 2-step Biotin stamps was slightly lower, while the 3xBiotin-7T stamp had the lowest correlation coefficient, corresponding to the poorest performance in reproducing the desired pattern.

**Figure 4.**
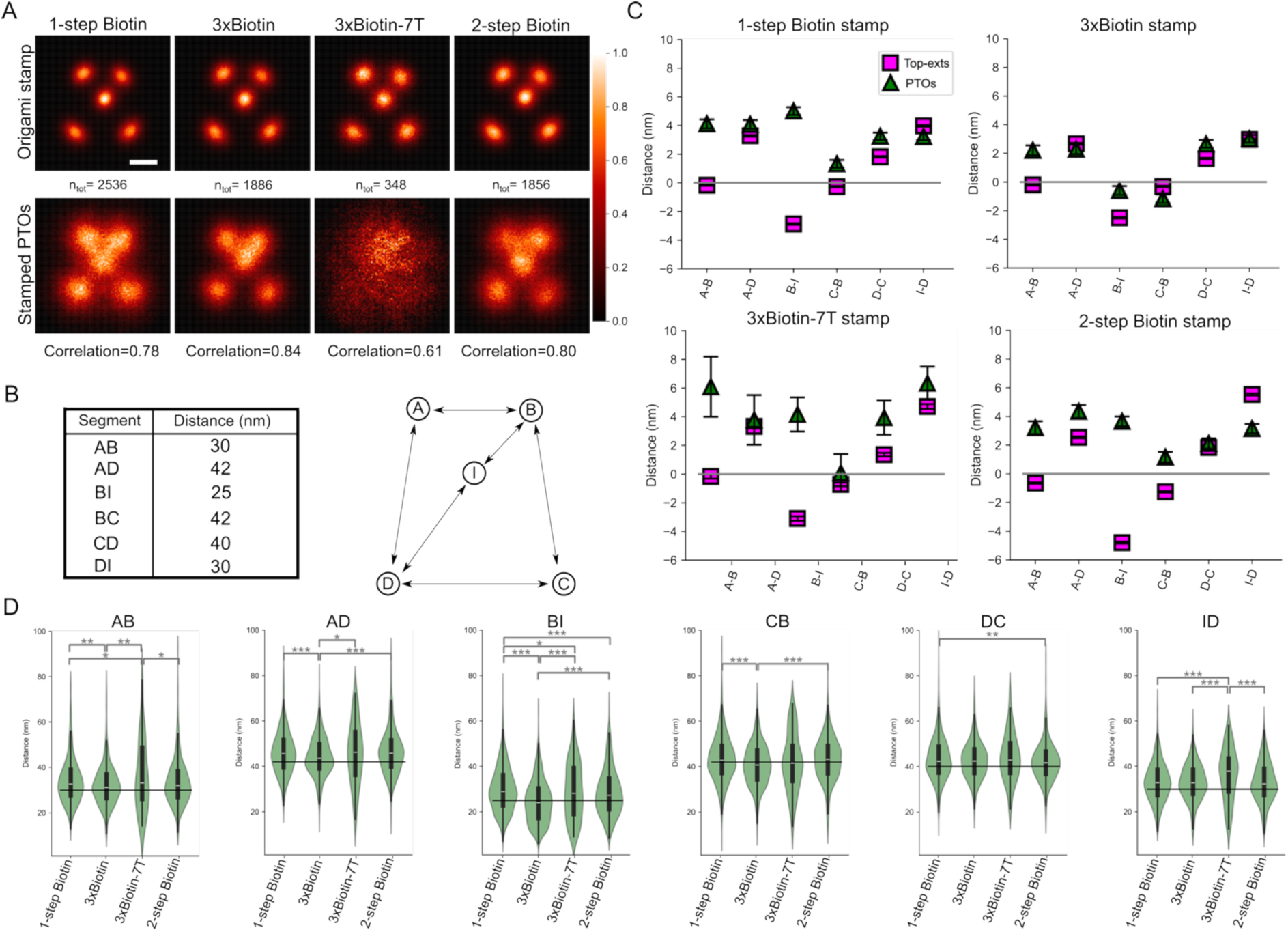
Stamping precision. (A) Heatmaps depicting the totality of the observed patterns, either constituted by Top-ext (Origami stamps) or stamped PTOs for each type of origami stamp. Each image was normalized to their maximum value, to visually compare the different origami stamp/stamped PTOs pairs. Scale bar 20 nm. The stamping precision was estimated by measuring the Pearson correlation coefficient between the two patterns (indicated below the images for each stamp). (B) The table lists the measurements of the designed distances estimated for the origami structure. On the right, the graphic representation of the pattern illustrates the position of each segment. (C) The deviation (in nm) of the measured Top-ext or PTO segment length from the designed distance length was obtained by subtracting the mean observed distances from the designed lengths (listed in B). The resulting values are depicted in the graphs and listed in Table S3. Error bars show the standard error of the mean (SEM) distance of Top-ext or PTOs while the grey line represents the deviation = 0 nm. (D) Distance distributions of the stamped segments for each origami stamp. Differences among distributions were statistically tested using a two-sided Kolmogorov-Smirnov test. The significance levels are * p<0.05, ** p<0.01=, *** p<0.001, and **** p<0.0001. Interquartile ranges (IQR) and median values are represented by black box plots and white lines, respectively. The black line represents the designed distance. The presented results are based on one representative experiment for each type of origami stamp.

While maintaining the geometry of the designed pattern is crucial, identifying ligand positions and their exact spacing could be instrumental in studies of multivalent ligand binding events. The characterization of the distance that separates them could potentially provide important molecular insights into the affinity and specificity of molecular interactions.

Distance (segment) measurements between the 5 different pattern elements, namely AB, AD, BI, CB, DC, and ID, were estimated from the origami design and reported in the table in Figure 4B. The distances between the stamped PTOs as well as those among the Top-exts of the origami stamp could deviate from the estimated measures due to the eventual structural distortion of the origami and the inherent localization precision (Table S1) of DNA-PAINT imaging. Therefore, we calculated the deviation (in nm) by subtracting the mean observed distances between Top-exts or PTOs from the designed distances across the four origami structures. The values obtained for each origami stamp system are reported in Table S3 and Figure 4C. The reference line at 0 nm represents no deviation from the designed segment lengths.

The maximum deviation of the position of the Top-ext segments observed across all the origami stamps is 5.5 nm (Table S3). Most of the measured deviations fell within the reported NeNA values (Table S1), suggesting that the observed deviations could be an intrinsic technical consequence of the localization precision achieved. The Top-ext segments of the 1-step, 2-step, and 3xBiotin-7T stamps presented similar but broader deviations (Figure 4C), ranging between 0.2 and 5.5 nm (Table S3). The shortest deviation range from the designed segment length was observed among the Top-exts of the 3xBiotin (Figure 4C), ranging only between 0.2 and 3.0 nm (Table S3), which falls well within the measured NeNA value.

Considering the overall deviation values and the localization precision (Table S1), the stamped patterns tended to exceed the expected distances of the segments. The 3xBiotin origami demonstrated the lowest deviation values, sfsignimilar to those recorded for the Top-ext counterpart (Figure 4C and Table S3), suggesting that this type of stamp was able to achieve a greater stamping precision than the other analyzed structures. The improved precision could be attributed to the branched PEG linker (∼3.5–4.0 nm), which increased the likelihood of one or more biotins reaching the nearest binding sites in an area perpendicular to the PTO projection.

Noticeably, the pattern stamped by the 3xBiotin-7T origami recorded most of the longest deviations with a range between 3.8 and 6.3 nm (Table S3). This was due to the overall broader distribution observed for the length of the segments between the stamped 3xBiotin-7T PTOs (Figure 4D), which was found to be highly different from the distributions of the other three origami stamps, particularly for distances equal to or shorter than 30 nm (AB, BI, and ID).

Therefore, adding a 7T linker (approximately 2.3 nm long) negatively affected the stamping precision. The prolongation of the 5′ ends of PTOs, summed to the length of the PEG linkers, might have dramatically deviated the binding to further neutravidin molecules, causing the highest variability of the inter-distance measurements of stamped PTOs. Additionally, the significant stamping deviation may have biased the distance-based cluster classification, introducing further imprecision in the distance measurements. For example, if two localization clusters belonging to separate PTOs are too close to each other, they may share a common pattern position. As a result, both clusters could be mistakenly classified as belonging to the same class. This represents a practical compromise, which our approach overcomes through the sheer sample size.

Finally, while the 1-step assembly improved the transfer efficiency by increasing the number of annealed PTOs, it was not able to achieve a precision of stamping similar to that of 3xBiotin origami. This result could be due to structural tension building up at the 4T single-stranded portion of the Bott-ext, protruding from the bottom surface of the origami. When annealed to the PTO, the Bott-ext-PTO complex might not be exactly perpendicular to the origami plane, leading to mild systematic distortion of the pattern.

## Conclusion

In this work, we demonstrated a versatile and accessible surface patterning approach using DNA origami as a stamp for oligonucleotide pattern transfer. By integrating the SMLM technique DNA-PAINT in the stamping process, we enabled single-molecule visualization of the origami stamps and performed rigorous nanoscale analysis of the stamped PTOs on the patterned surface. Our flexible “bottom-up” approach eliminates the need for expensive, specialized equipment typically required in “top-down” methods. It also enables adaptation to different surfaces by modifying the 5’ ends of PTOs with various reactive groups for covalent and non-covalent immobilization. The use of a 2-step assembly process enables easy exchange of PTOs, facilitating the patterning of a wide range of ligands across diverse surfaces and conditions. We further developed a robust method for the identification and automated quantification analysis of individual stamp-stamped pattern pairs, offering quantitative insights into patterning efficiency and precision. While asymmetrical patterns proved more effective in preventing rotational misalignments, the method remains adaptable to any pattern that can be resolved using DNA-PAINT or related SMLM techniques.

Looking forward, this approach could be adapted for applications where the strong negative charge of DNA origami negatively affects the activity of the binder under investigation. Additionally, if residual DNA from the PTO interferes with the system, DNase treatment can be applied. In such scenario, the ligand can be attached either to the 5′ end of the biotinylated nucleotide or to upstream nucleotides, potentially preventing further digestion downstream of the functionalized site.

Our stamping technique opens new avenues for studying nanoscale distance-dependent biological processes such as molecular interactions in receptor activation,^40^ multivalent binding,^41^ engineering enzymatic cascades,^42^ or the activity of molecular motors ^43^ across extended spatial scales, and various detection techniques. While different stamp designs showed varying effectiveness for specific applications, future work could explore larger scale geometries by placing origami structures at a fixed orientation ^44^ or building 2D lattices of origami tiles.^45^ Overall, this method represents a significant advance in accessible and versatile surface patterning strategies for biological studies.

## Methods

### Origami assembly and characterization

The origami structure design was based on M13mp18 ssDNA scaffold using the CaDNAno software^46^, inspired by the work of Ke and colleagues^47^. Staple strand extensions, the PTO, and the invader sequences (Suppl. Tables S4 and S5) were purchased from Integrated DNA Technologies (IDT). The M13mp18 scaffold and staples strands were diluted to a concentration of respectively 10 nM and 100 nM in 1xTAE buffer (10x, Invitrogen), supplemented with 10 mM of MgCl_2_. The PTO was diluted to a concentration of 5 µM for the 1-step assembly of the extended origami. The assembly program consisted of incubation at 65°C for 10 minutes (min) followed by a gradient from 63°C to 55°C with a decrease of 0.1°C every 5 min, a gradient from 55°C to 47°C decreasing of 0.1°C every 4 min, a gradient from 47°C to 40°C decreasing of 0.1°C every 3 min, a gradient from 40°C to 25°C decreasing of 1° every min, ending with a final storage temperature of 10°C. The assembled DNA origami was purified using a 100kDa cutoff Amicon Ultra centrifugal filter (Merck Millipore). The spin filter was equilibrated with TAEM buffer (1xTAE, 10 mM MgCl_2_) at a speed of 2000 g for 5 min. The origami was diluted to 500 μl, added to the filter, and spun for 5 min at 2000 g. This last step was repeated seven times, then the origami was collected and quantified by Nanodrop (DeNovix). In the 2-step assembly process, Biotin-PTOs were added in 10-fold excess respect to the purified extended origami structure, and the mix was incubated at 30°C for 1 hour (h). Excess Biotin-PTOs was removed in a second purification step as previously described. To assess the presence of annealed Biotin-PTOs, 20-folds excess of neutravidin (Sigma Aldrich) was added to the assembled origami and incubated at room temperature (RT) for 30 min. The purity of the origami and its ability to bind neutravidin was assessed by agarose gel electrophoresis. The gel was prepared by adding 1% w/v agarose (Thermo Fischer Scientific) in 1xTBE buffer (10x, Invitrogen) supplemented with 10 mM MgCl_2_. Samples were run through the gel using 80 V, for 2.5 h.

For the TEM characterization of the origami stamps, grids (Electron microscopy Sciences, CF400-Cu) were glow-discharged (PELCO easyGlow) for 45 seconds (s), then 5 μl of a 5 nM origami was added to the grid and incubated for 1.5 min. Unbound origami were removed followed by the sequential addition of 4 μl of negative staining buffer (2% urinyl formate in 20 mM KOH). The first application was quickly removed while the second application was incubated for 20 s. Upon staining, the grid was air-dried and imaged by using FEI Tecnai G2 Electron Transmission microscope equipped with a TVIS CMOS camera (TemCam-F416, 4k). The acquired data were finally averaged and aligned using the Scipion software^48^.

### PTO conjugation to Biotin

HPLC-purified Biotin-PTO was purchased from IDT along with the PTO and the 7T-PTO sequences bearing a 5′ NH_2_ group. To obtain the 3xBiotin-PTO and the 3xBiotin-7T-PTOs, the 5′ NH_2_ moiety was reacted with 10-fold excess NHS-PEG5-tris-PEG4-alkyne (Conjuprobe) in 100 mM 4-(2-hydroxyethyl)-1-piperazineethanesulfonic acid (HEPES) pH 8 and 60 % dimethyl sulfoxide (DMSO), overnight (O/N) at 17°C and 500 rpm. The reaction was followed by ethanol precipitation by adding 300 mM CH_3_COONa pH 5.2, 1 μg/μl of glycogen, and 5x volume of cold absolute ethanol to each reaction. The mix was centrifuged at 17.000 g for 30 min, the supernatant discarded, the pellet mixed with 200 μl of cold ethanol, and spun at the same speed for 10 min. This last step was repeated twice to remove the excess NHS-PEG5-tris-PEG4-alkyne, and the pellet was air-dried and resuspended in nuclease-free water.

To couple three azide-PEG3-biotin (Sigma-Aldrich) to PTO-/7T-PTO-PEG5-tris-PEG4-alkyne by Cu-catalyzed cycloaddition, a “click buffer” was prepared by mixing 5.33 µl of 5 mM CuSO4, 1.07 µl of 50 mM Tris(benzyltriazolylmethyl)amine (TBTA), 5 µl of 200 mM ascorbate and into a final volume of 43.4 vol% DMSO. The reaction mix was prepared by adding 1/3 of the total reaction volume of click buffer, 35% DMSO, 100mM HEPES pH 7.4, the PTO or 7T-PTO bearing the branched linker, and 30x excess azide-PEG3-biotin. The reaction was carried out at 27°C, 700 rpm for 4.5 h. The mix was diluted in water and the conjugate was purified by Reversed-Phase HPLC (RP-HPLC). Fractions containing the conjugate were freeze-dried and resuspended in water. Quality control was carried out through 16% denaturing PAGE (17 ml UreaGel Diluent (National Diagnostics), 19 ml UreaGel Concentrate (National Diagnostics), 4 ml 10x TBE (Gibco by Life Technologies), 420 μl ammonium persulfate (APS) (Sigma Aldrich), and 20 μl TEMED (Sigma-Aldrich)).

### PEG-b-neu-coated surfaces preparation

Poly(acryl-amide)-g-(PEG-N_3_,1,6-hexanediamine,3-aminopropyldimethylethoxysilane) (3500:116.2:161.3 Mr; 0.15:0.425:0.425) PAcrAm-g-PEG-N_3_(NH2,Si) were synthesized and characterized by SuSoS, Switzerland^33^. A modified version of this polymer with terminal Biotin, PAcrAm-g-PEG-biotin(NH_2_, Si)^35^, was used for the surface preparation to achieve an ultradense antifouling PEGylated surface covered with neutravidin for the pattern stamping process. To prepare the PEG-b-neu surfaces, borosilicate glass coverslips were cleaned by ultrasonication in acetone and isopropanol alcohol and subsequently dried under a stream of N_2_. To activate the surface and remove residual organic contamination, the coverslips were treated with an oxygen plasma reactive ion etching for 3 min (Vision 300 MK II, Advanced Vacuum) with a radio frequency generator power of 100 W, 100 SCCM O_2_ flow at 25 mTorr pressure. Subsequently, the coverslips were placed on a droplet of 0.1 mg/ml PAcrAm-g-PEG-biotin(NH_2_, Si) in 1 mM HEPES buffer at pH 7.4 and incubated for 30 min, followed by rinsing for 30 s in deionized water and N_2_ drying. To maximize neutravidin coating of the surface and the following immobilization of biotinylated molecules, without negatively affecting the anti-fouling properties^35^, the substrate was exposed to a 0.2 µm filtered ceric ammonium nitrate-based chromium etchant solution (Sigma-Aldrich) for 30 s^34^. The surface was rinsed with deionized water and reincubated with PAcrAm-g-PEG-N_3_(NH_2_, Si) at 0.1 mg/mL in similar fashion before the final rinse with deionized water, and drying under a stream of N_2_ before being mounted on an ibidi sticky-Slide VI 0.4.

### Surface Plasmon Resonance

Biacore SIA Au chips (GE Healthcare) were sputter-coated with 4 nm Ti, followed by 30 nm SiO_2_ to resemble the glass coverslips’ surface chemistry. Subsequently, the surface was PEGylated and prepared for functionalization as described above. The neutravidin and DNA origami were respectively diluted to 100 µg/ml and 6 nM in buffer C (1x PBS + 500 mM NaCl, 1mM EDTA, 0.05% Tween-20), and injected on the surface for 20 min. The invader strand was injected at a concentration of 300 nM. The buffers and solutions were degassed by sonication and filtered with a 0.22 µm syringe filter prior to use. A Biacore 3000 was used for the measurements. All the experiments were performed at a flow rate of 5 µl/min at 25°C.

### Molecular stamping and TIRFM imaging

The PAcrAm-g-PEG-biotin surface supplied as a 6-channel Ibidi slide was further functionalized with 100 µg/ml of neutravidin diluted in 1x PBS and incubated for 20 min at RT. The surface was washed three times with 1x PBS and three times with buffer B+ (10 mM MgCl_2_, 5 mM Tris-HCl pH 8, 1 mM EDTA, 0.05% Tween-20, pH 8). The origami stamps were added to the surface at a concentration of 0.5 nM and incubated at RT for 15 min. Unbound structures were removed by washing three times with buffer B+ and three times with buffer C. Gold nanoparticles (80 nm, Sigma Aldrich) were diluted 1:4 in buffer C and incubated on the surface at RT for 5 min. Unbound particles were removed by washing with buffer C. The channel of interest was mounted on a TIRF inverted microscope (Oxford Nanoimager S) equipped with a sCMOS camera with a pixel size of 117 nm, a 100x oil-immersion objective with a numerical aperture (NA) of 1.4, and a built-in autofocus system. At each imaging session, the relative Cy3B-labeled DNA imager sequence was injected into the channel in a volume of 130 μl at a concentration of 1 nM diluted in buffer C supplemented with 2.5 mM protocatechuic acid (PCA), 10 nM protocatechuate-3,4-dioxygenase (PCD), and 1 mM 6-hydroxy-2,5,7,8-tetramethylchroman-2-carboxylic acid (Trolox). Buffer C was used to wash the imagers away, followed by the injection of the invader strand at a concentration of 300 nM diluted in TAEM supplemented with 50 mM NaCl and incubated for 15 min at the temperature reached on the microscope stage (28-29°C). The surface was then thoroughly washed with buffer C to remove excess invaders and the origami in solution, and the second imager was added to visualize the stamped PTOs. Fluorescent imaging was performed in TIRF mode using a 561 nm laser line and a laser power of 20 mW, as reported by the acquisition software NimOS. A total number of 20000 frames were recorded for each acquisition, with an exposure time of 100 ms to obtain a field of view of 428x684 pixels (approximately 50x80 μm).

### Data treatment and estimation of stamping efficiency and precision

Raw fluorescence data were processed for reconstruction by the Picasso software package^30^ adapted to GB-sized TIFF files. Picasso Localize was used to localize and fit (LQ, Gaussian method) the single blinks in every acquired frame. The resulting processed HDF5 file was opened in Picasso Render and the rendered 2D image was initially drift-corrected by using the redundant cross-correlation (RCC) process and successively, using the “pick” tool, gold nanoparticles were manually picked as fiducial markers for a second round of drift-correction. The resulting drift-corrected data was then filtered using Picasso Filter by plotting sx and sy coordinates in a 2D plot. Filtered rendered data of the Top-exts of the origami stamps and their relative stamped PTOs were both opened in Picasso Render and aligned using several rounds of RCC and picked gold nanoparticles. NeNA values were automatically calculated from Picasso Render software.

To identify localizations originating from the Top-exts for individual well-assembled stamps, origami patterns were identified in Picasso render by drawing a ROI enclosing the structure (Figure S7 A). While 40-50 stamps were manually picked, the remaining structures in the FOV were automatically segmented by the “pick similar” tool based on the manually picked localizations. The localizations from all picked Top-exts and the associated PTO were further analyzed using the following custom-made python script: <u>anon.erda.au.dk/share_redirect/afbdOYXXO5</u>.

Here, both the origami stamp and the corresponding stamped patterns were rotationally aligned and refined using only the Top-exts as references for the determination of the angle of rotation (Figure S7 B). Due to the high degree of symmetry of the designed pattern, the rotational alignment was performed by determining the angle of rotation for the maximum Pearson correlation in relation to a computationally simulated reference. This resulted in a primary ensemble of the Top-ext signal. By using the simulated reference, within-ROI clusters were then classified as points A, B, C, D, or I using a distance-based classifier, identifying the most probable point of origin for the localizations (Figure S7 C). Following this classification, the distances between centroids in the Top-exts patterns were obtained (Figure S7 D) for assessment of any 90° rotational errors still present in the primary ensemble.

To quantify the number of rotational alignment errors, the distances between the Top-exts were analyzed using a two-component Gaussian mixture model. The determined models are shown in Figure S8 and Table S6. The patterns were analyzed according to their rotational classes determined from the models (Figure S7 D), and filtered to remove patterns containing rotational errors. The patterns correctly rotated were collected into the final ensembles, and the intra-cluster distances between Top-ext or PTOs were measured (Figure S7 E). The distance distributions calculated from the ensemble ROIs are presented in the violin plots in Figure S9.

### Statistical analysis

The distribution of measured distances presented in Figure 4D were statistically tested using a two-sided Kolmogorov-Smirnov test. Significance levels are * p<0.05, ** p<0.01=, *** p<0.001, and **** p<0.0001.

## Acknowledgments

This work was supported by the Danish National Research Foundation (CellPAT, DNRF135). The work was in part funded by the Novo Nordisk Foundation (RNA-META, NNF23OC0081177, and NNF21OC0071574). Mette Galsgaard Malle is funded by the Lundbeck Foundation (R380-2021-1393). We wish to thank Rasmus Schøler Sørensen for his guidance on the design of the origami structure and Juliàn Valero Moreno for his advice on the design of the stamping mechanism. We would like to acknowledge Kasper Okholm for his contribution to the preliminary assessment and optimization of the functionalized surfaces and Lasse Hyldgaard Klausen for AFM imaging of previous origami designs.

## Author contributions

L.T. conceptualization, validation, microscopy imaging and data treatment, manuscript writing, editing, and data visualization. A.S. conceptualization, surface preparation, SPR, imaging optimization, and manuscript editing. E.A.S. analysis and statistics of DNA-PAINT data, data visualization, and manuscript editing. A.S. and E.A.S. contributed equally to the work. X.Z. DNA origami structure design and characterization by TEM. M.G.M supervision (E.A.S), DNA-PAINT analysis and statistics, manuscript editing, and data visualization. D.S.S. conceptualization, supervision, and manuscript editing. J.K. conceptualization, supervision, and manuscript editing.

## Supplemental information

**Nanoscale precise stamping of biomolecule patterns using DNA origami**

Laura Teodori, Ali Shahrokhtash, Elisabeth Asta Sørensen, Xialin Zhang, Mette Galsgaard Malle, Duncan Stewart Sutherland, and Jørgen Kjems.

## Supplemental Figures

**Figure S1.**
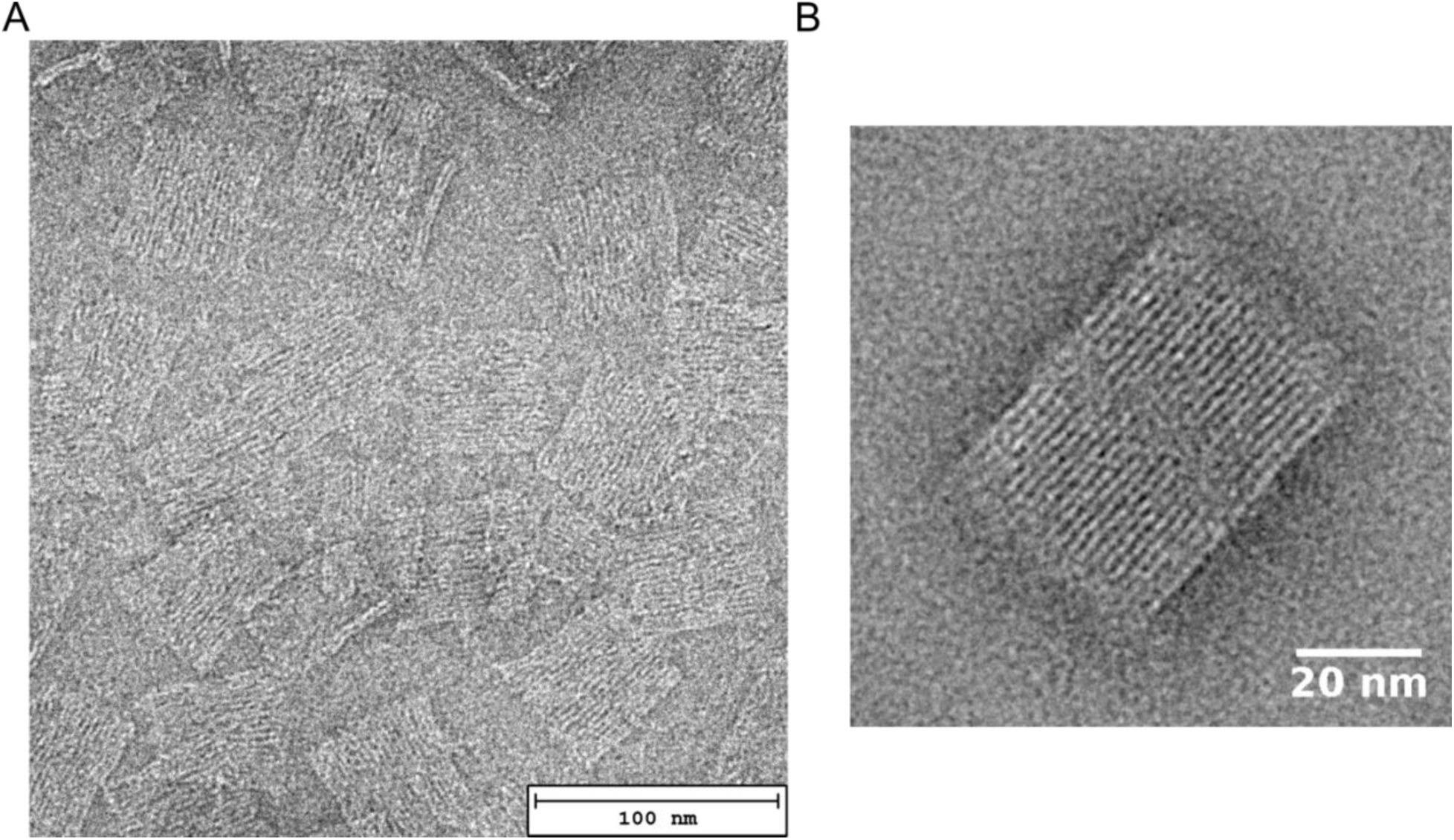
TEM images of the origami stamp. (A) Raw image of origami stamps on a grid. (B) Scipion software 2D rendering of the origami stamp upon averaging of over 100 manually selected structures.

**Figure S2.**
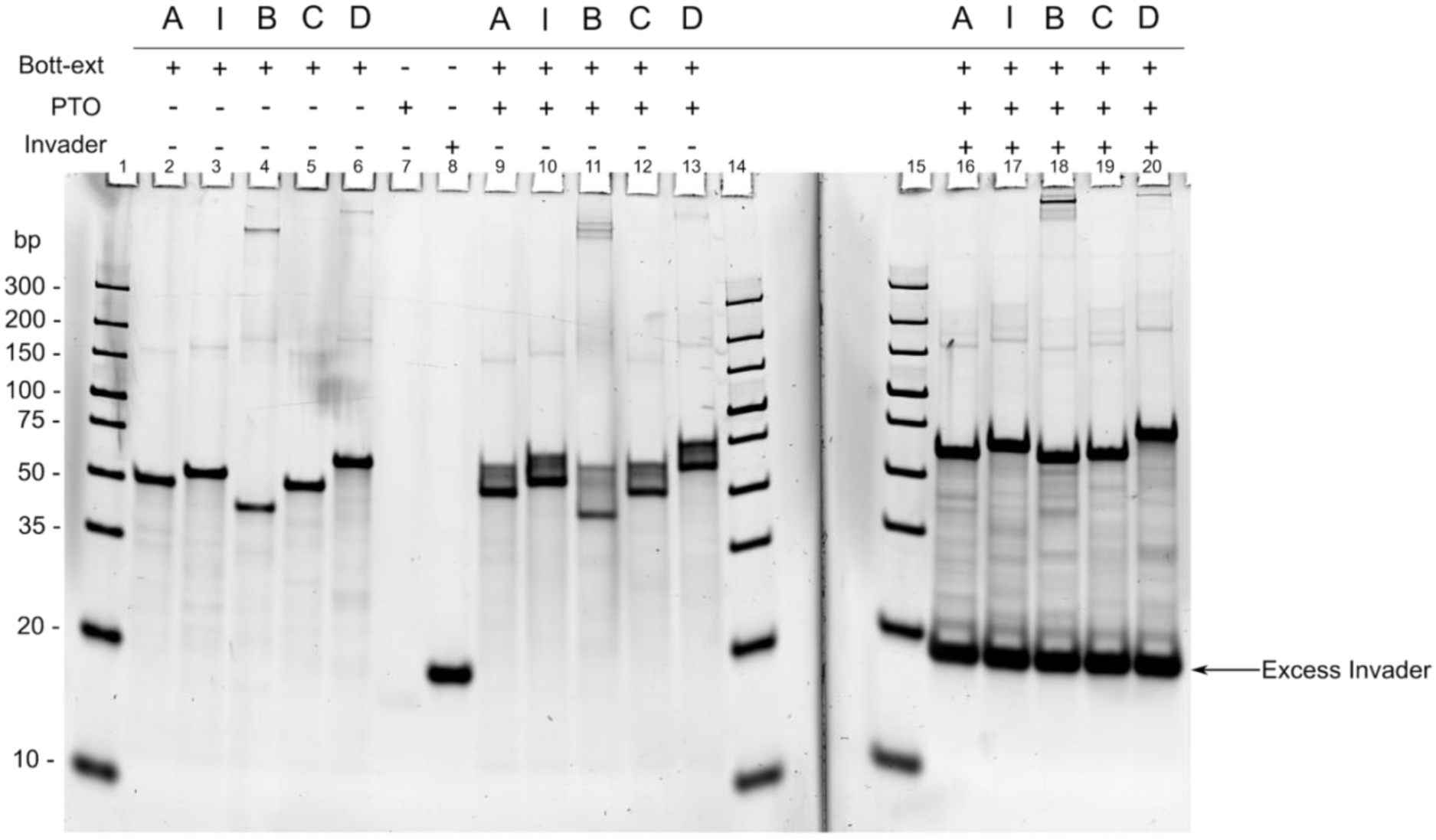
Toehold-mediated strand displacement mechanism. 10% acrylamide native gel showing Bott-ext oligonucleotides (wells No. 2-6), biotinylated PTO (well No. 7), and the invader sequence (well No. 8). PTO were annealed to each single Bott-exts and the complexes were visualized in the same gel (wells No. 9-13) as an upward shift. Part of the Bott-ext-PTO complexes were mixed with excess invader sequence and loaded in the gel (wells No. 16-20). A larger upward shift was observed in this case, given the longer length of the invader oligonucleotide, which displaced the PTO and annealed to the Bott-exts from their 3’ toehold. The gel was stained with SYBR gold stain. Unfortunately, the single PTO and the strand-displaced PTOs (wells No. 16-20) were poorly visible or invisible, as the PTO is not efficiently stained by SYBR Gold.

**Figure S3.**
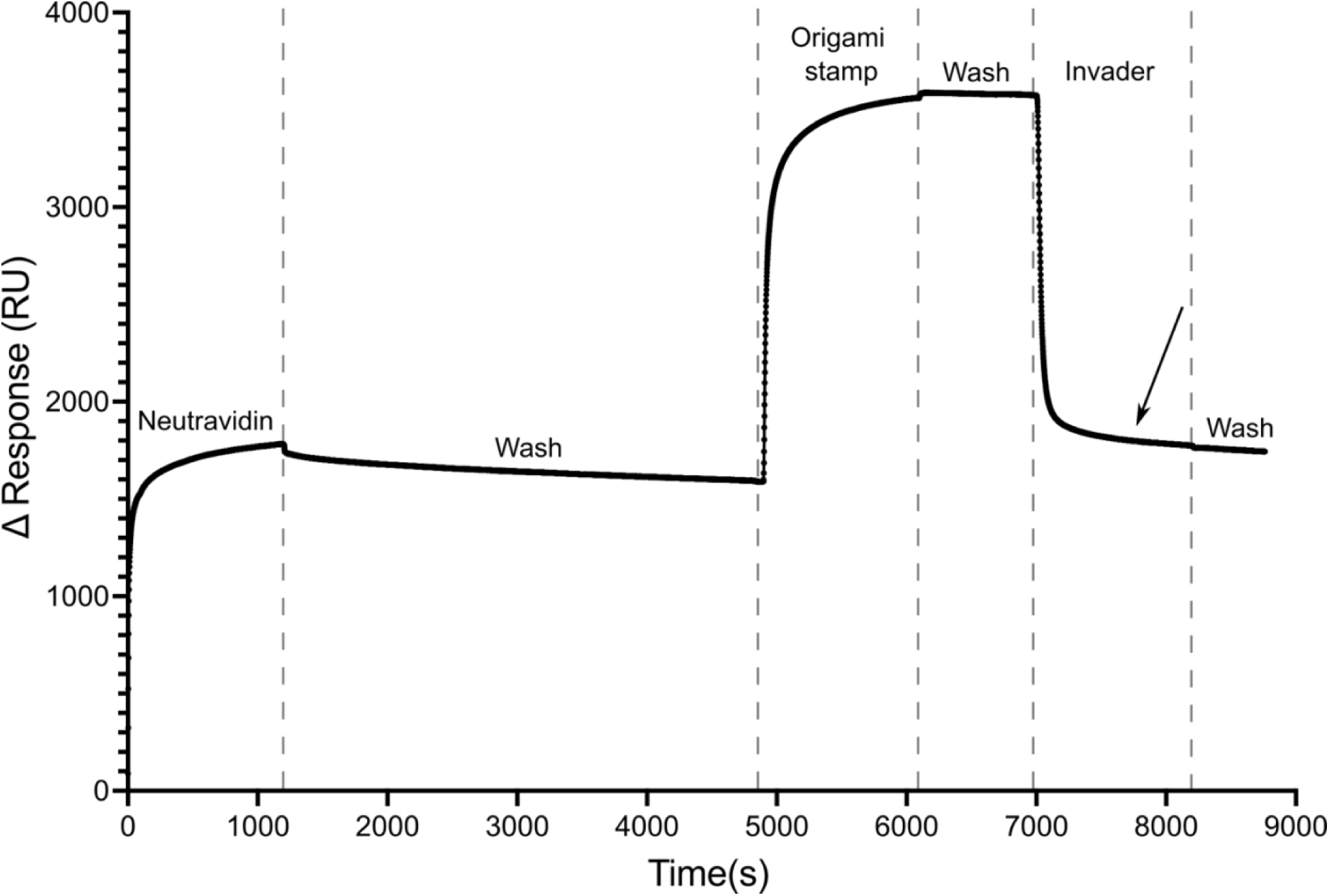
SPR-monitored stamping mechanism. Molecular stamping mechanism characterization by SPR, where the PEG-b-neu surface was generated by coating the PEG-biotin brush with neutravidin (from 0 to 5000 s), followed by the immobilization of the origami (from 5000 to 7000 s), indicated by an increase in the response. The application of the invader oligonucleotide (from 7000 s) caused a quick decrease in the response, indicating the release of the origami structures in solution. The final response (black arrow) indicated the presence of residual signal after the invasion, suggesting the presence of bound stamped PTOs, as a result of the complete stamping process.

**Figure S4.**
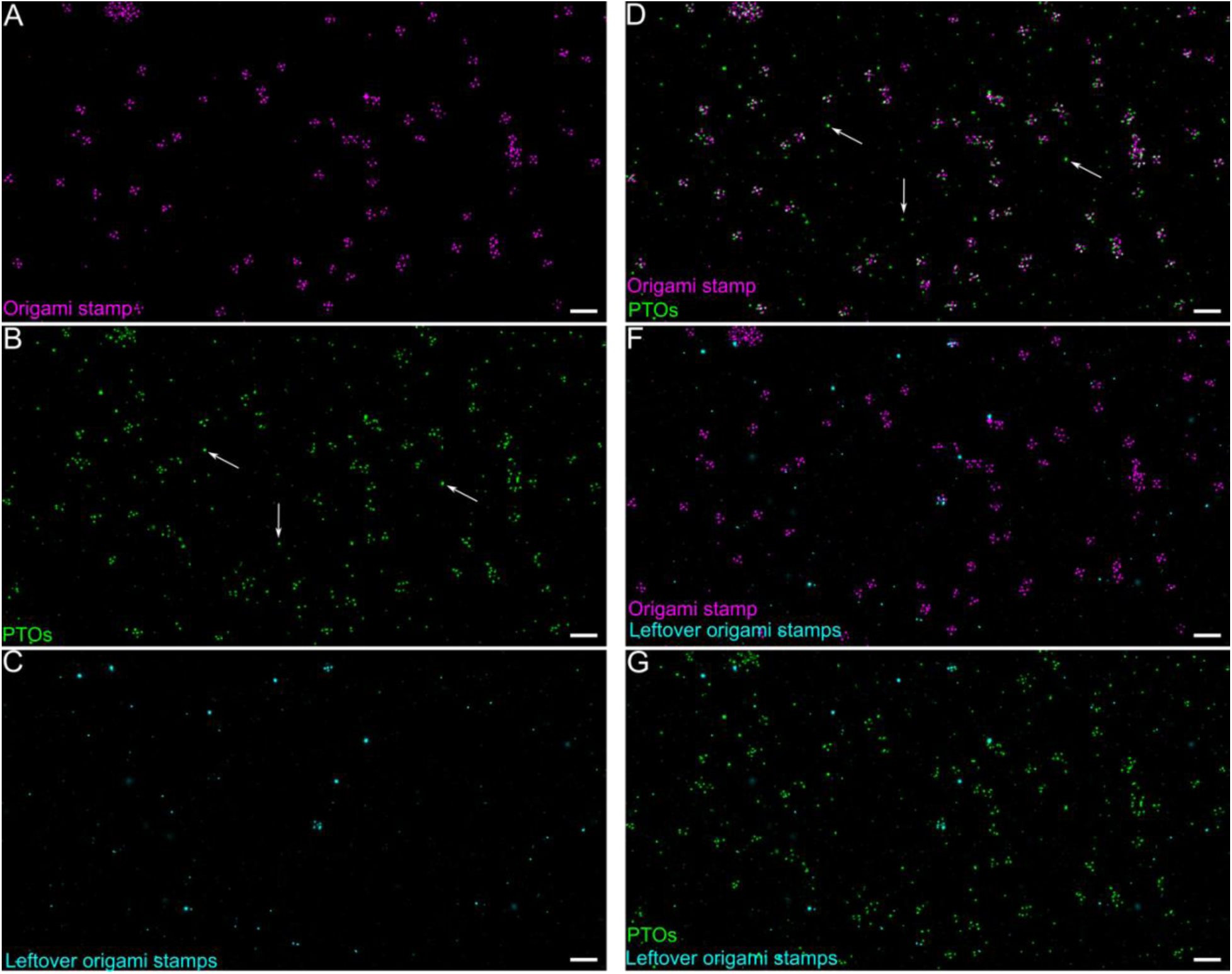
Signal of non-filtered origami stamps and single-stranded PTOs in magnified FOV. (A) Non-filtered signal of origami stamps; (B) Non-filtered signal of single-stranded PTOs. White arrows indicate PTO signals unrelated to stamped patterns; (C) Non-filtered signal of origami stamps, visualized after the stamping mechanism, showing the leftover origami stamps not successfully released in solution during strand displacement; (D) Merged signals of origami stamps and single-stranded PTOs. White arrows indicate PTO signals unrelated to stamped patterns; (E) Merged signals of origami stamps and leftover Origami, after stamping, and (G) Merged signals of single-stranded PTOs and leftover origami. Scale bar 200 nm.

**Figure S5.**
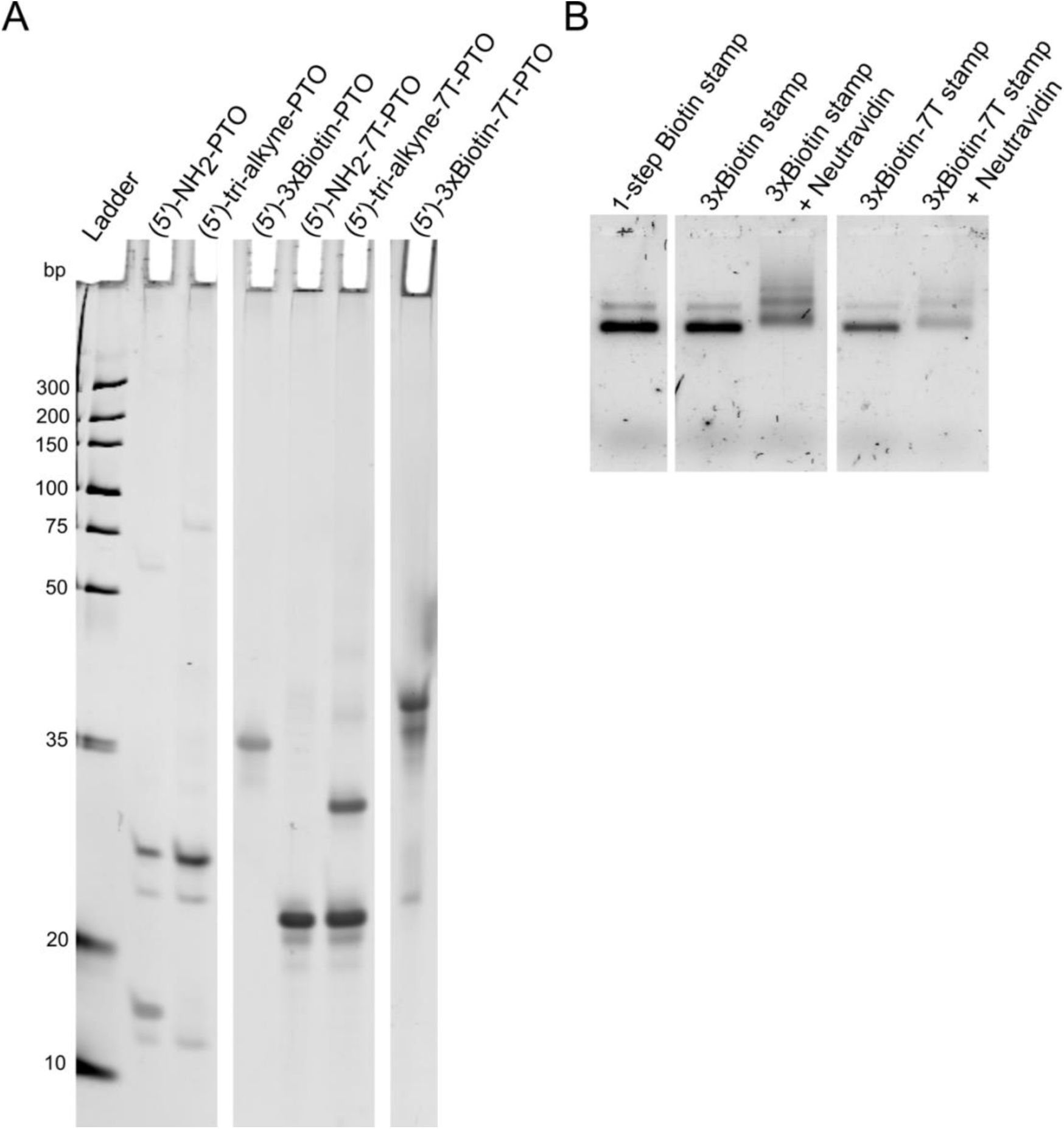
Conjugation of 3xBiotin-PTO and 3xBiotin-7T-PTO. (A) A 16% denaturing PAGE gel of 5′-NH_2_-PTO and 5′-NH_2_-7T-PTO conjugated to tri-alkyne linker and successively functionalized with three biotin molecules. (B) origami stamp assembled either with biotin-PTOs (1-step Biotin stamp), 3xBiotin-PTOs (3xBiotin stamp) and 3xBiotin-7T-PTOs (3xBiotin-7T stamp). The last two origami were incubated with excess neutravidin to verify by mobility shift assay the presence of biotinylated PTO annealed to the stamp.

**Figure S6.**
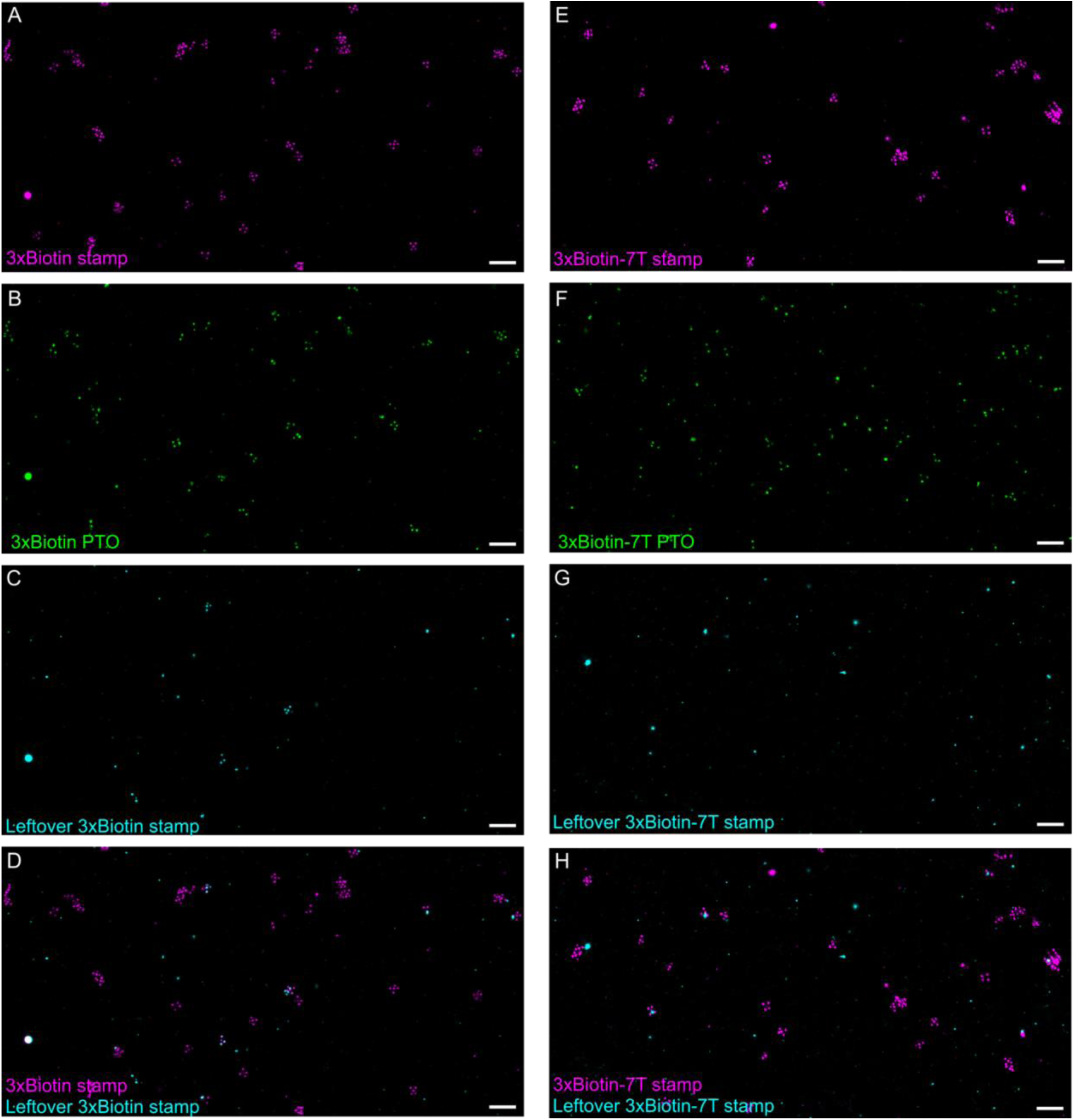
View of 3xBiotin and 3xBiotin-7T stamping. (A) Non-filtered signal of 3xBiotin stamps; (B) Non-filtered signal of 3xBiotin PTOs; (C) Non-filtered signal of leftover 3xBiotin stamps; (D) Merged signal of 3xBiotin stamps and leftover 3xBiotin stamps after strand displacement; (E) Non-filtered signal of 3xBiotin-7T stamps; (F) Non-filtered signal of 3xBiotin-7T PTOs; (F) Non-filtered signal of leftover 3xBiotin-7T stamps; (G) Merged signal of 3xBiotin-7T stamps and leftover 3xBiotin-7T stamps after strand displacement. Scale bar 200 nm.

**Figure S7.**
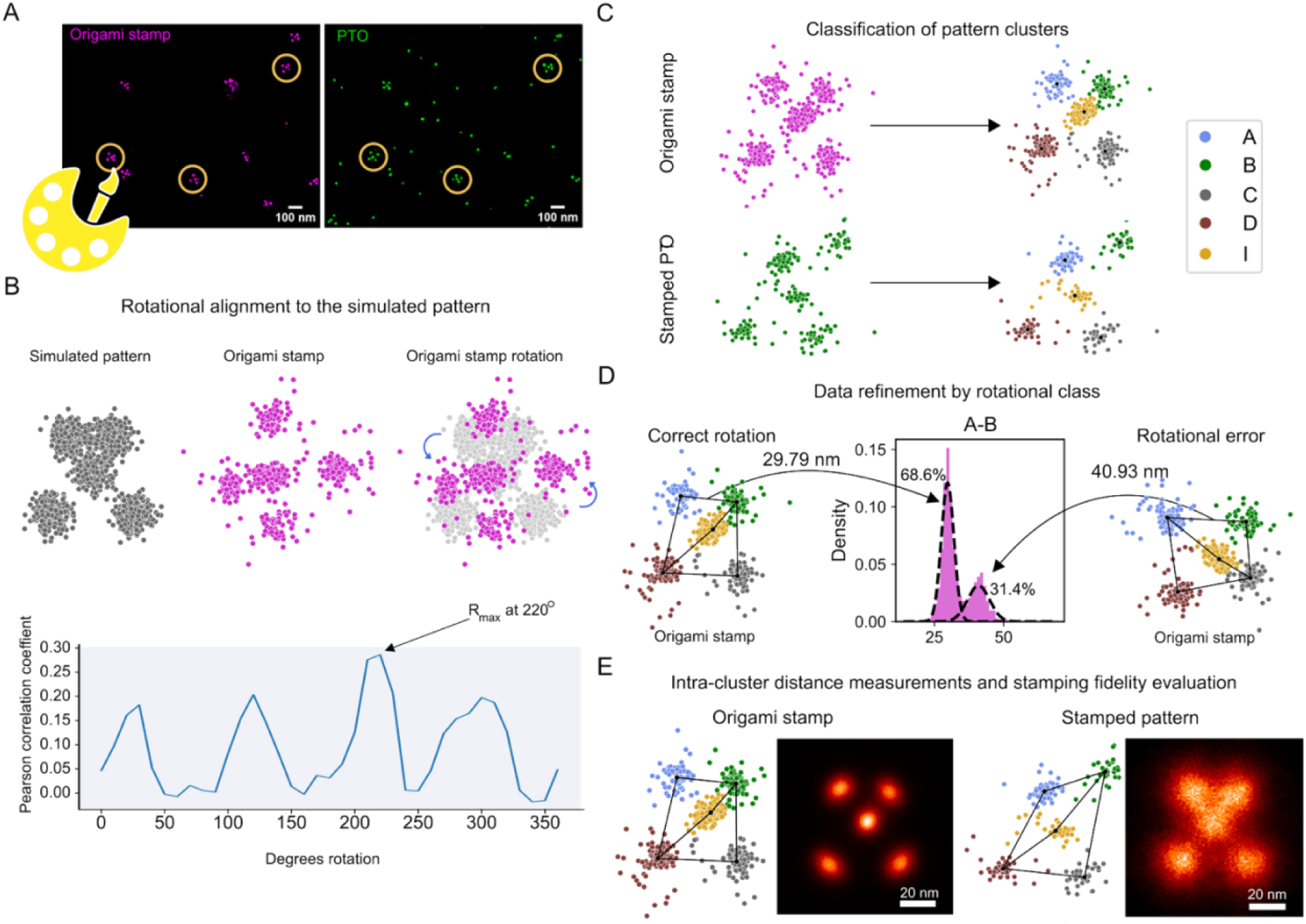
Data Processing. (A) Origami patterns were identified and selected in Picasso Render, and the corresponding localizations of the origami stamp and PTOs were exported for processing in the custom script. (B) The origami stamp localizations were rotated and aligned onto a simulated pattern. The angle of rotation was determined from the maximum Pearson correlation coefficient (bottom). (C) The aligned localizations were further classified using a simulated distance reference to assign the most probable strand of origin for each smaller cluster in each ROI. The cluster centroid is marked in black. (D) To identify rotational errors, the center of mass for each Top-ext cluster was used as the point of origin for the determination of intra-cluster distances. Each distance was analyzed using a two-component Gaussian mixture model (Figure S8) to obtain rotational models and identify the set of erroneously rotated patterns. Patterns containing rotational errors were filtered out, and the remaining origami stamp-stamped pattern pairs were used in the final ensemble. (E) Intra-cluster distances were measured for all the Top-exts and PTOs pattern pairs to determine the stamping fidelity of each origami stamp.

**Figure S8.**
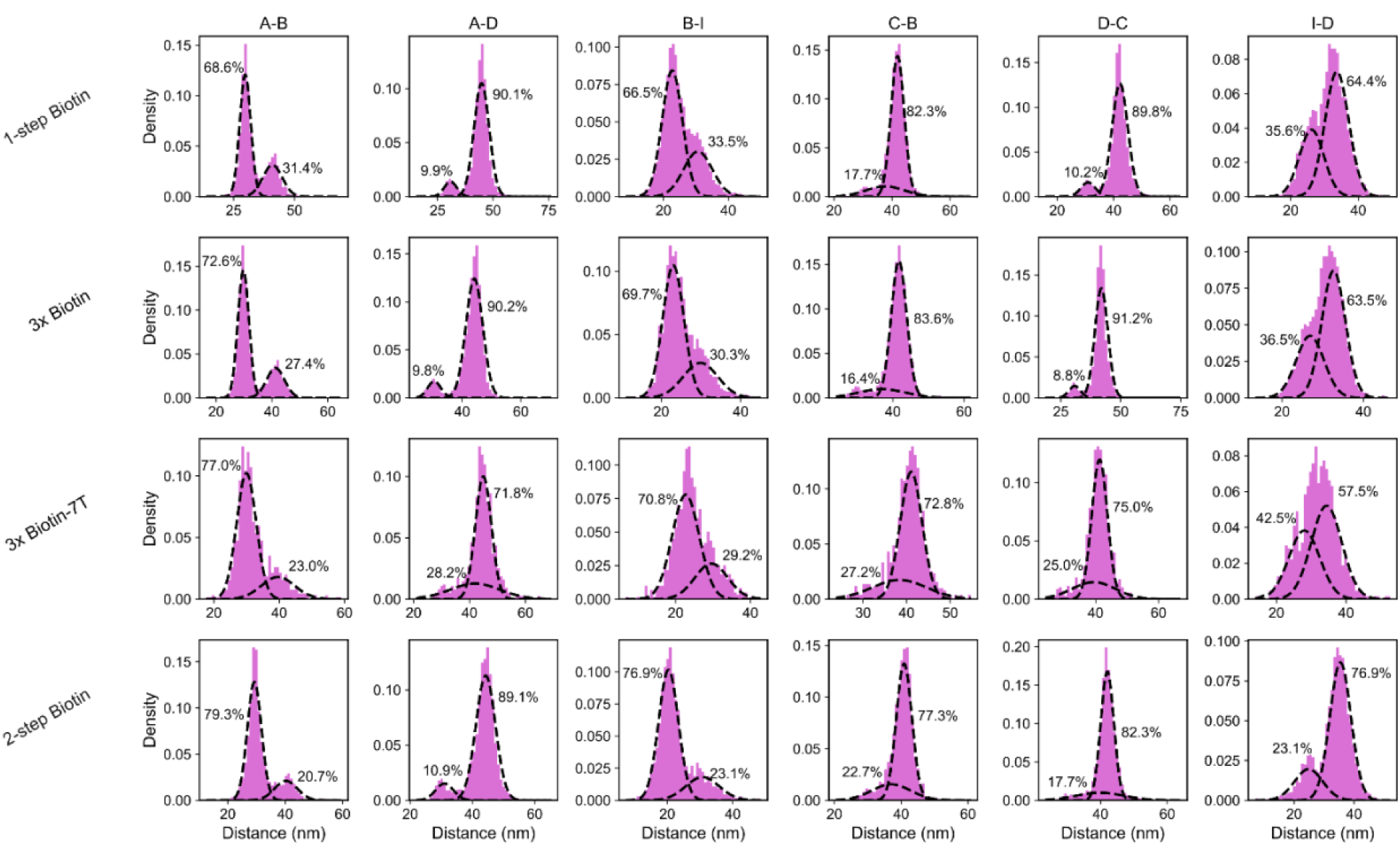
Gaussian mixture model decomposition of the distributions of measured distances after rotational alignment. Across each origami stamp, the measured distances between the observed Top-exts were analyzed using a two-component Gaussian mixture model. The weights are shown as percentages next to the respective component, and the parameters of the derived components are provided in Table S6. The resulting models were used to distinguish and filter erroneously rotated origami stamps and their corresponding PTO from the ensemble, as shown in Figure S7.

**Figure S9.**
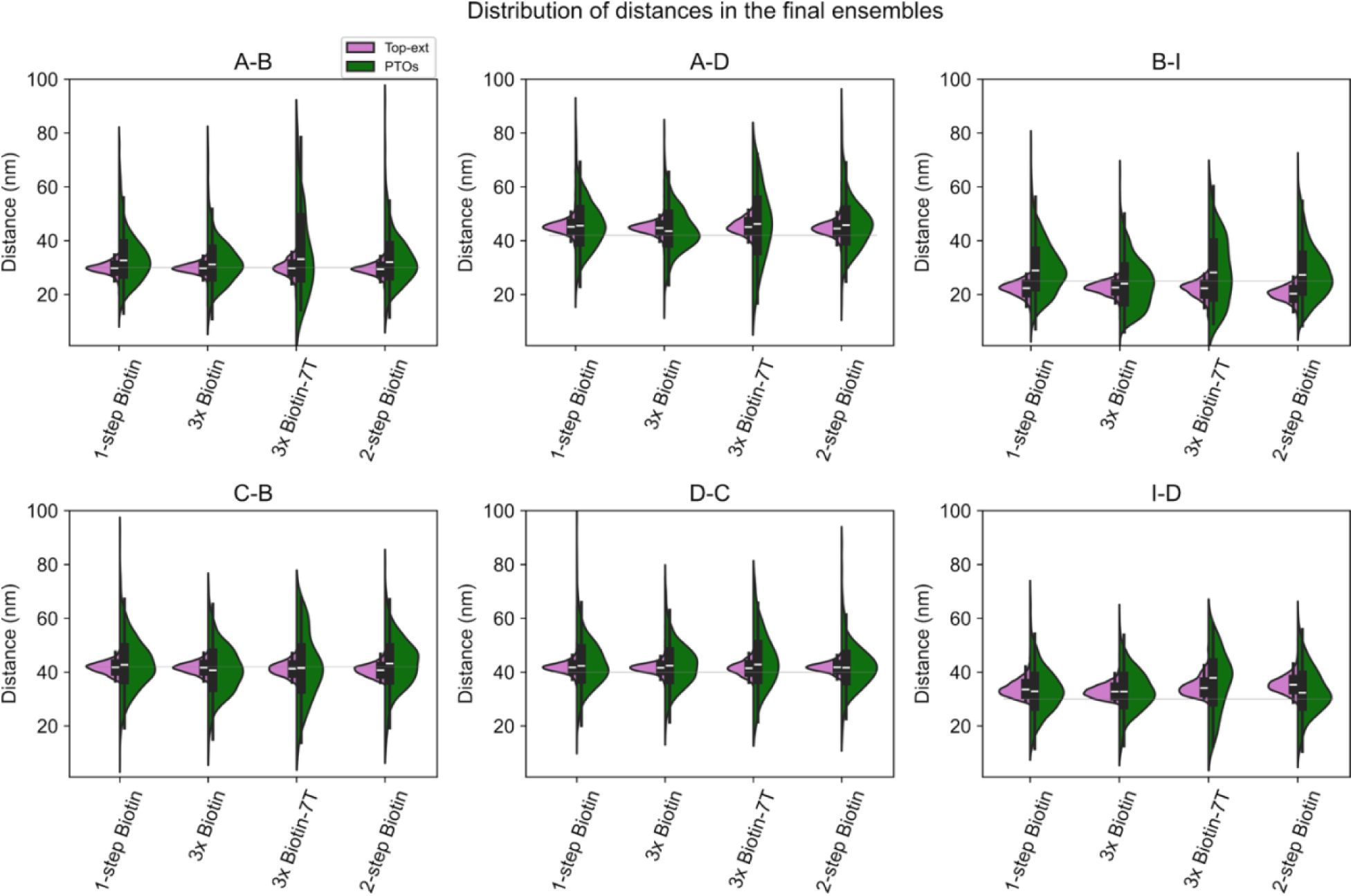
Filtered distributions of the measured distances. Violin plots and boxplots of the measured distances between Top-exts of the origami and PTOs after data filtration according to the rotational model in Figure S8. The white line and boxes show the median and IQR. The horizontal grey lines represent the designed distances.

## Supplemental Tables

**Table S1.**
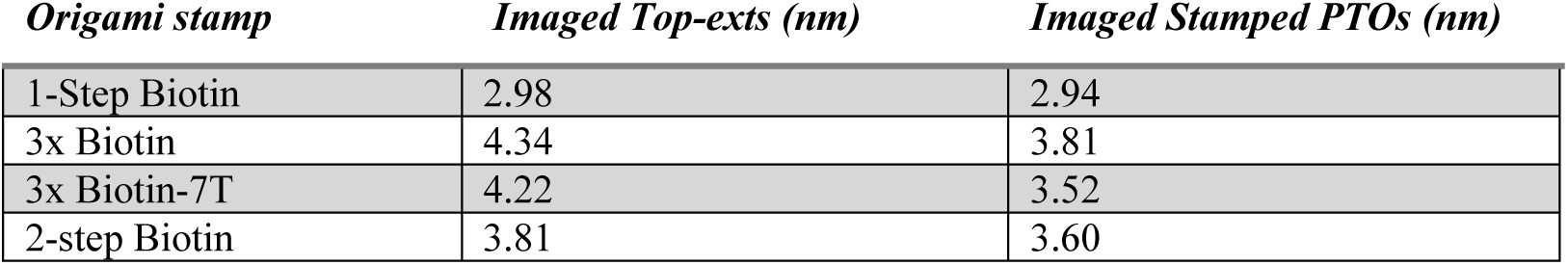
Averaged localization precision (nm) calculated by Nearest Neighbor Analysis (NeNA)

**Table S2.**
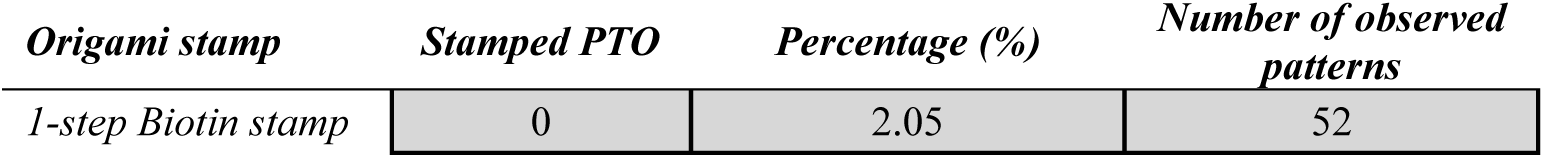
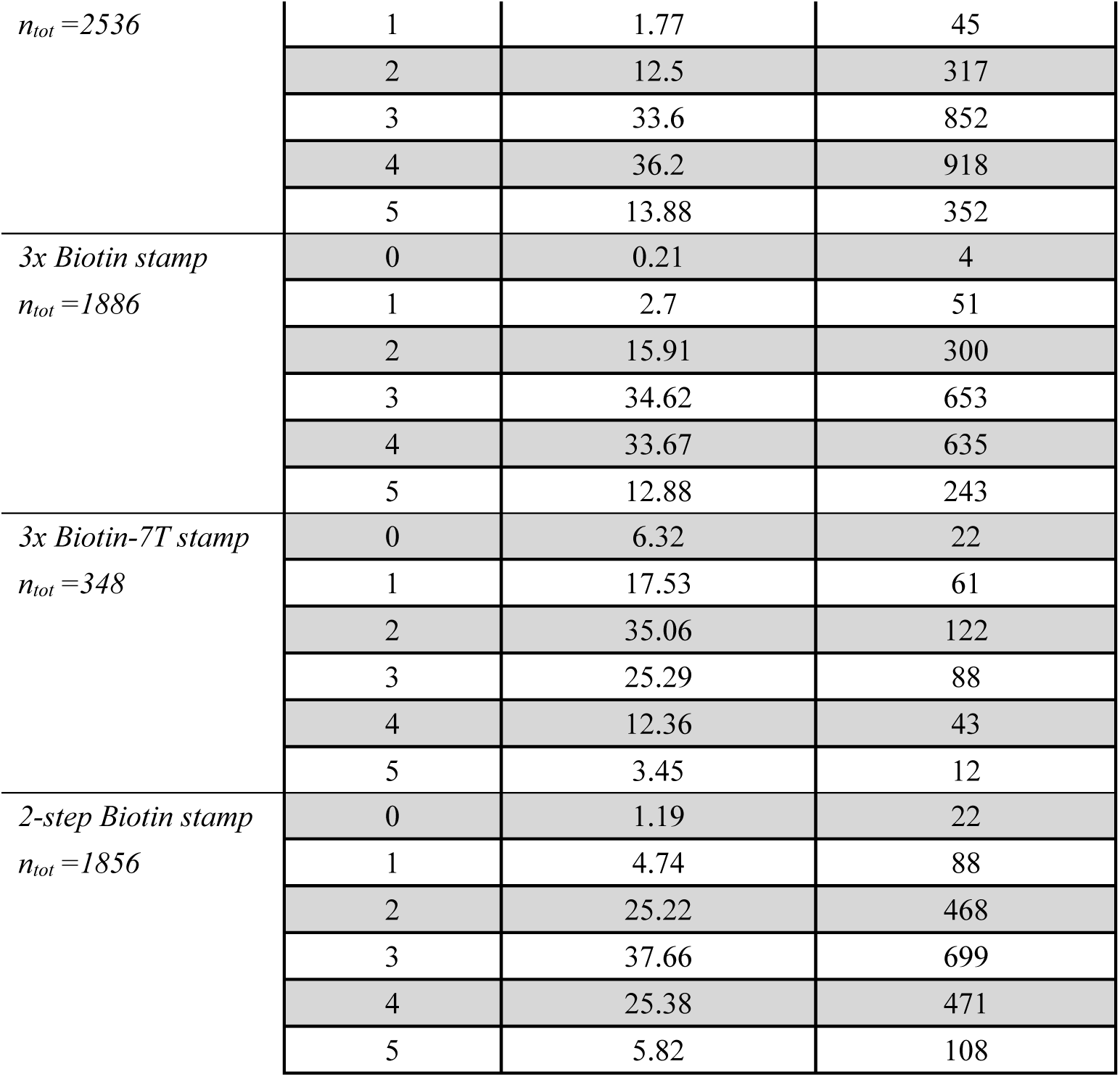
Percentage of fully, partially, and failed stamped patterns.

**Table S3:**
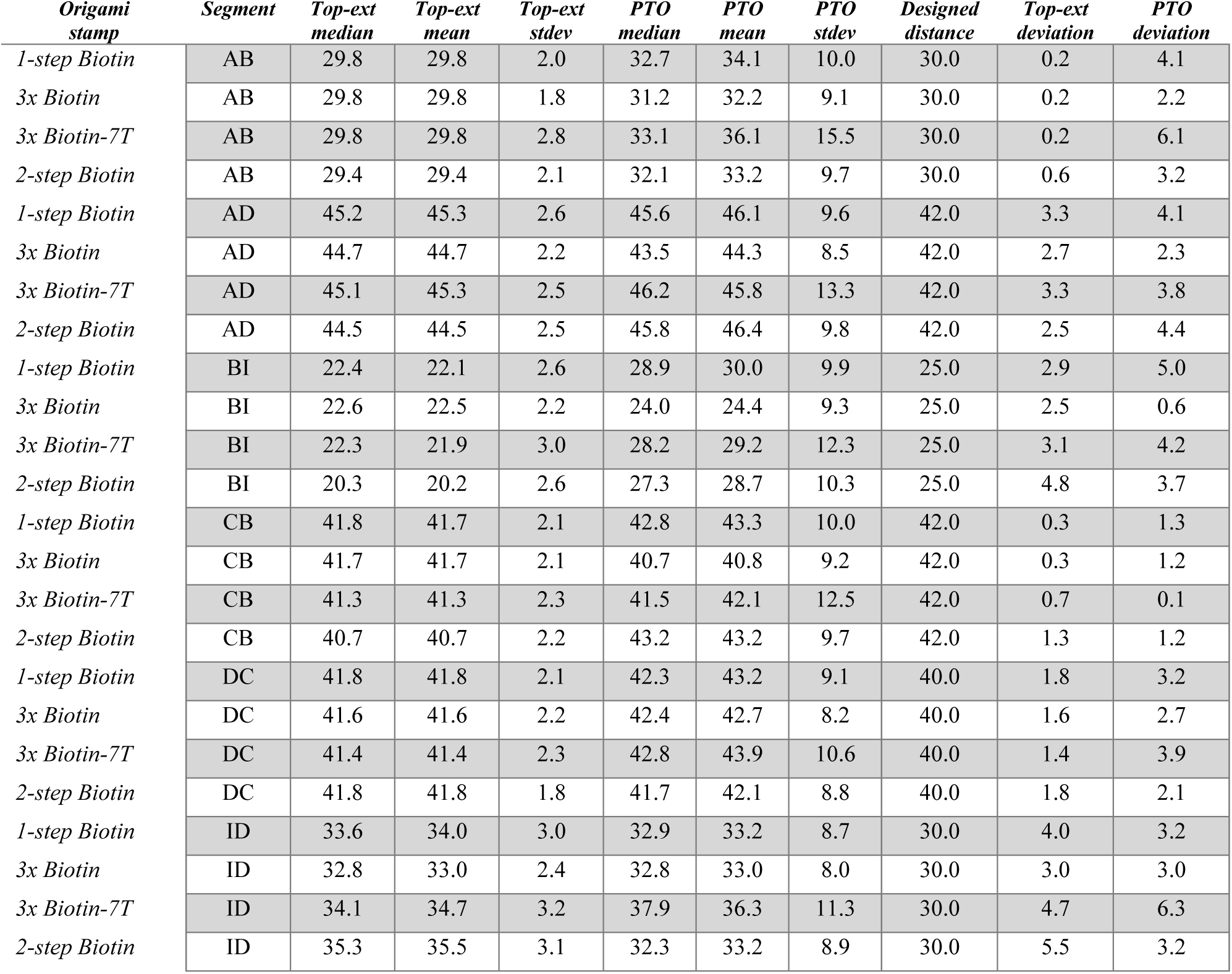
Summary statistics of the different segments and the deviation to the designed distances. The segments were measured across the different stamps and their stamped PTOs. The median, mean and standard deviations (stdev) values are shown to provide a reference of the physical scale of the stamping. The designed distances are used as references to obtain the absolute average distance deviation from the experimental mean values. All values are in nm.

**Table S4.**
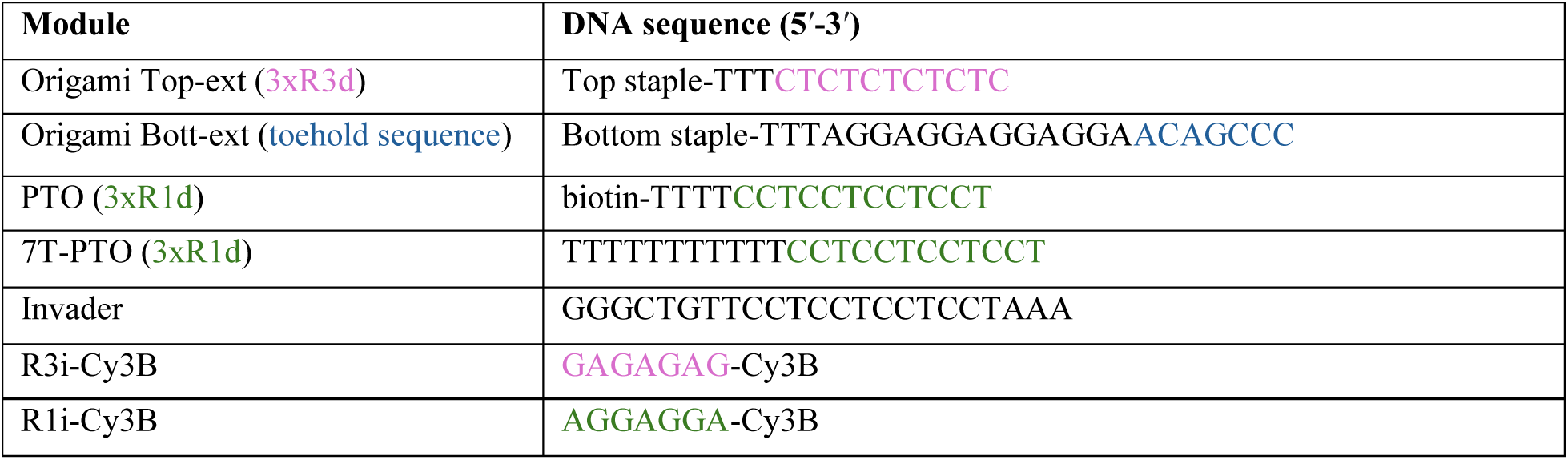
Sequences of staple extensions, PTOs, invader, and imager strands.

**Table S5.**
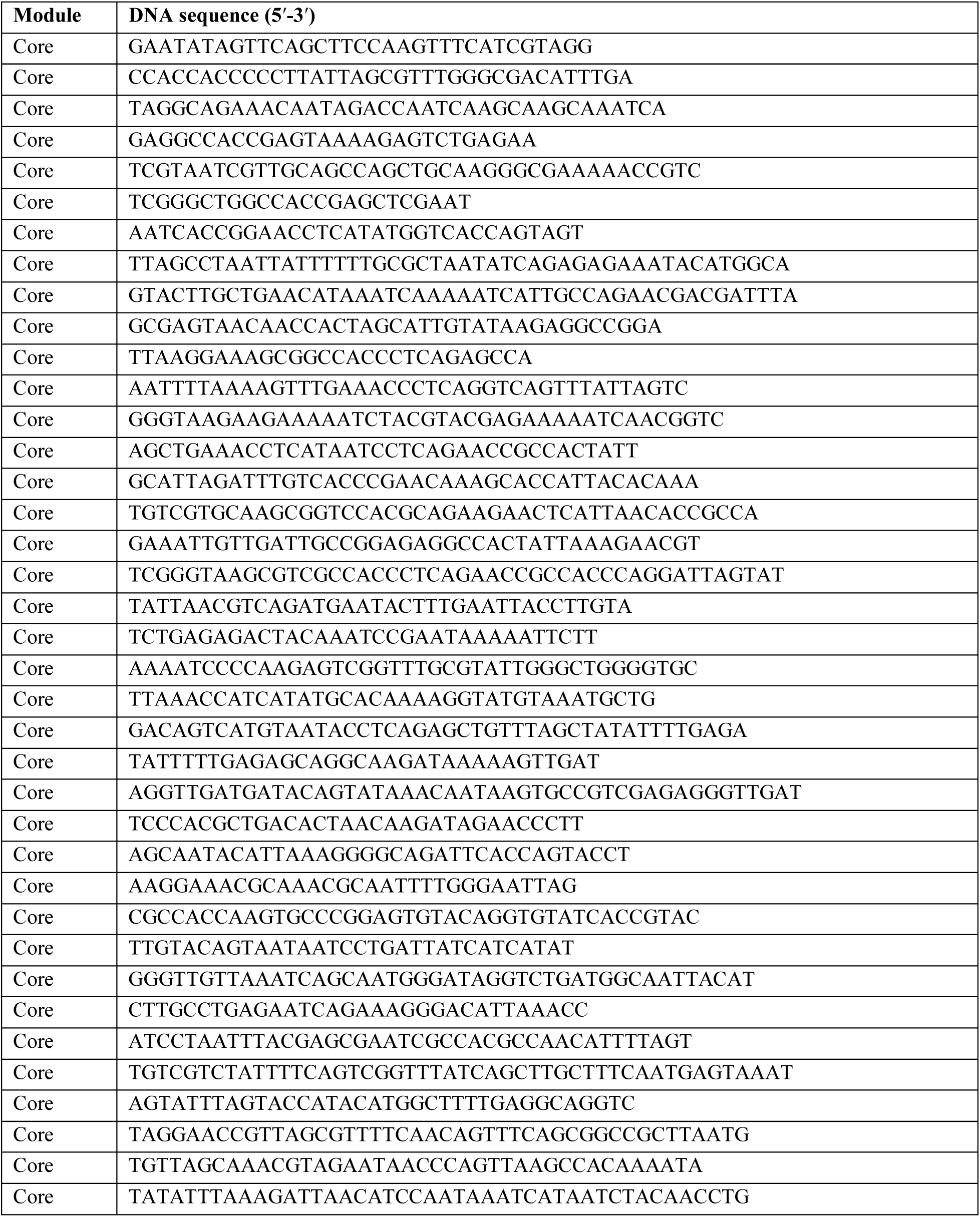
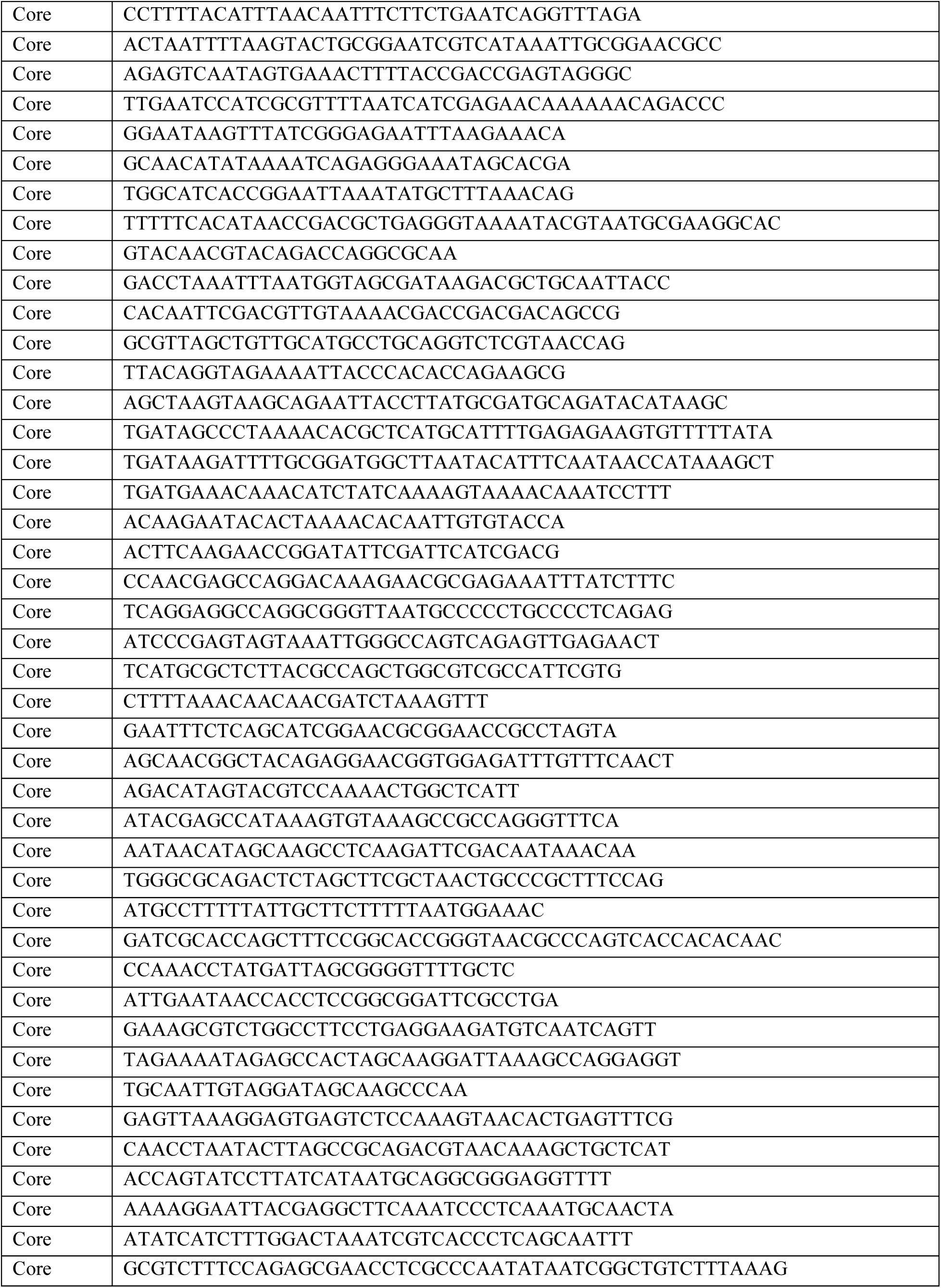
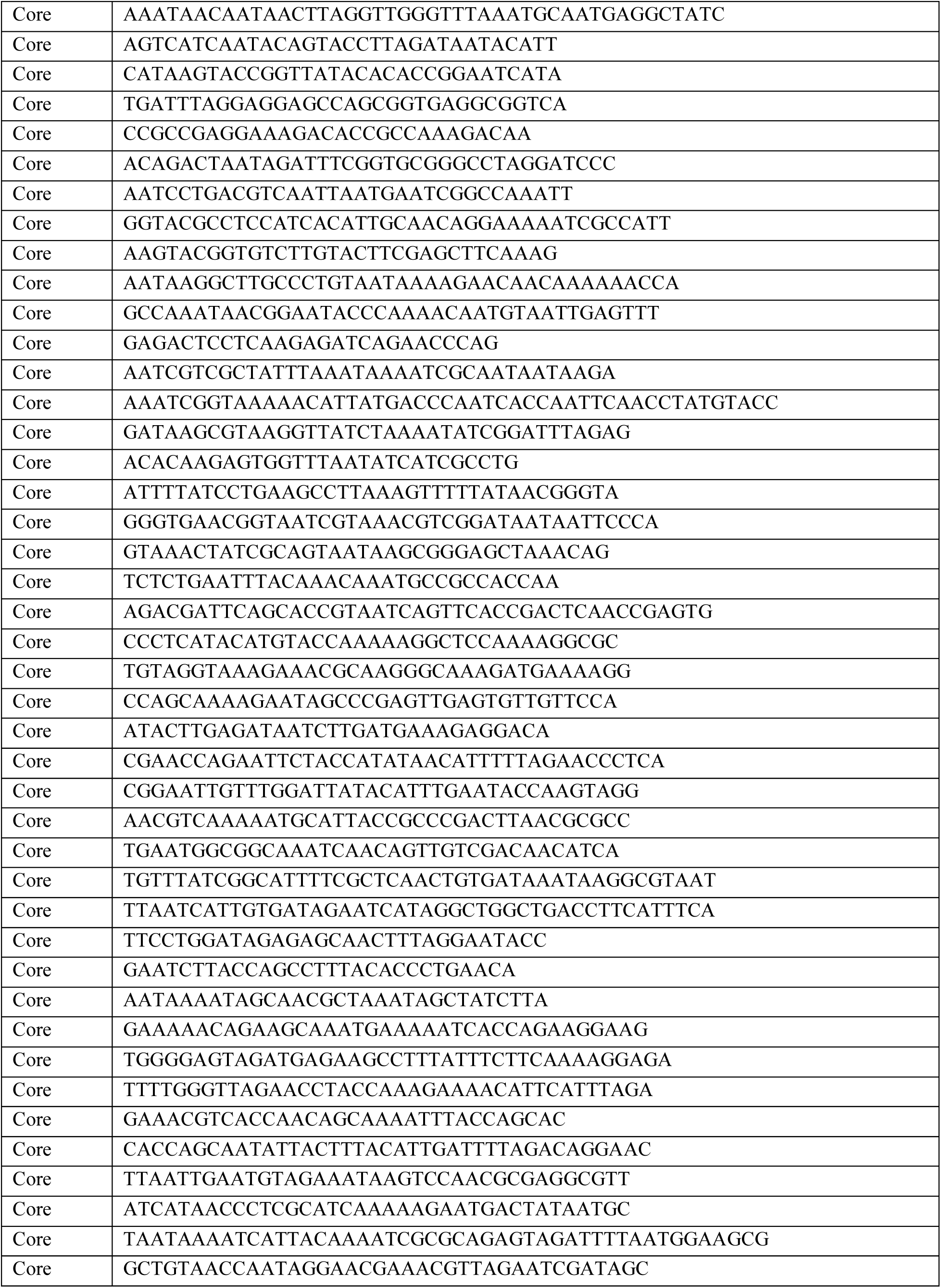
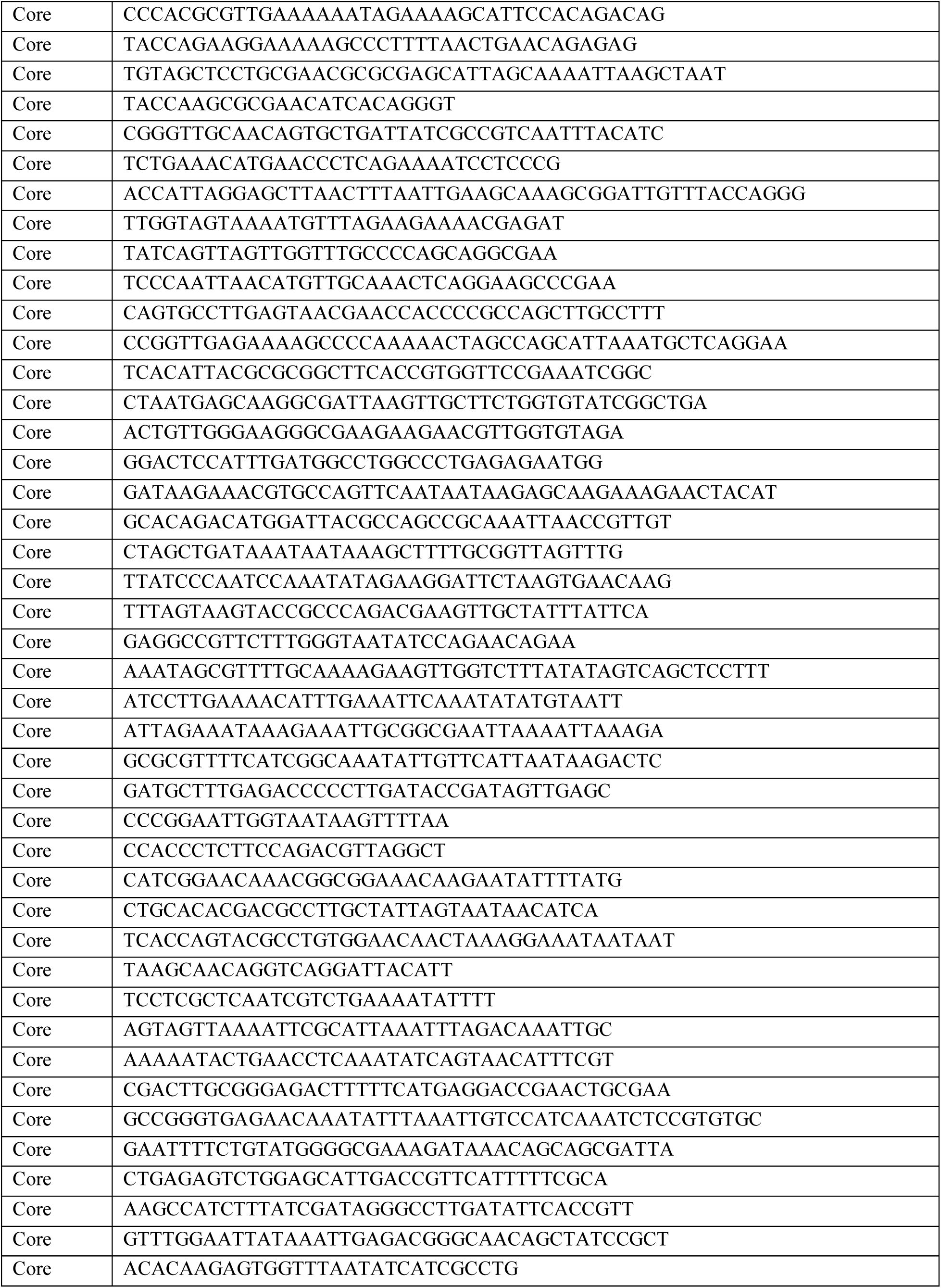
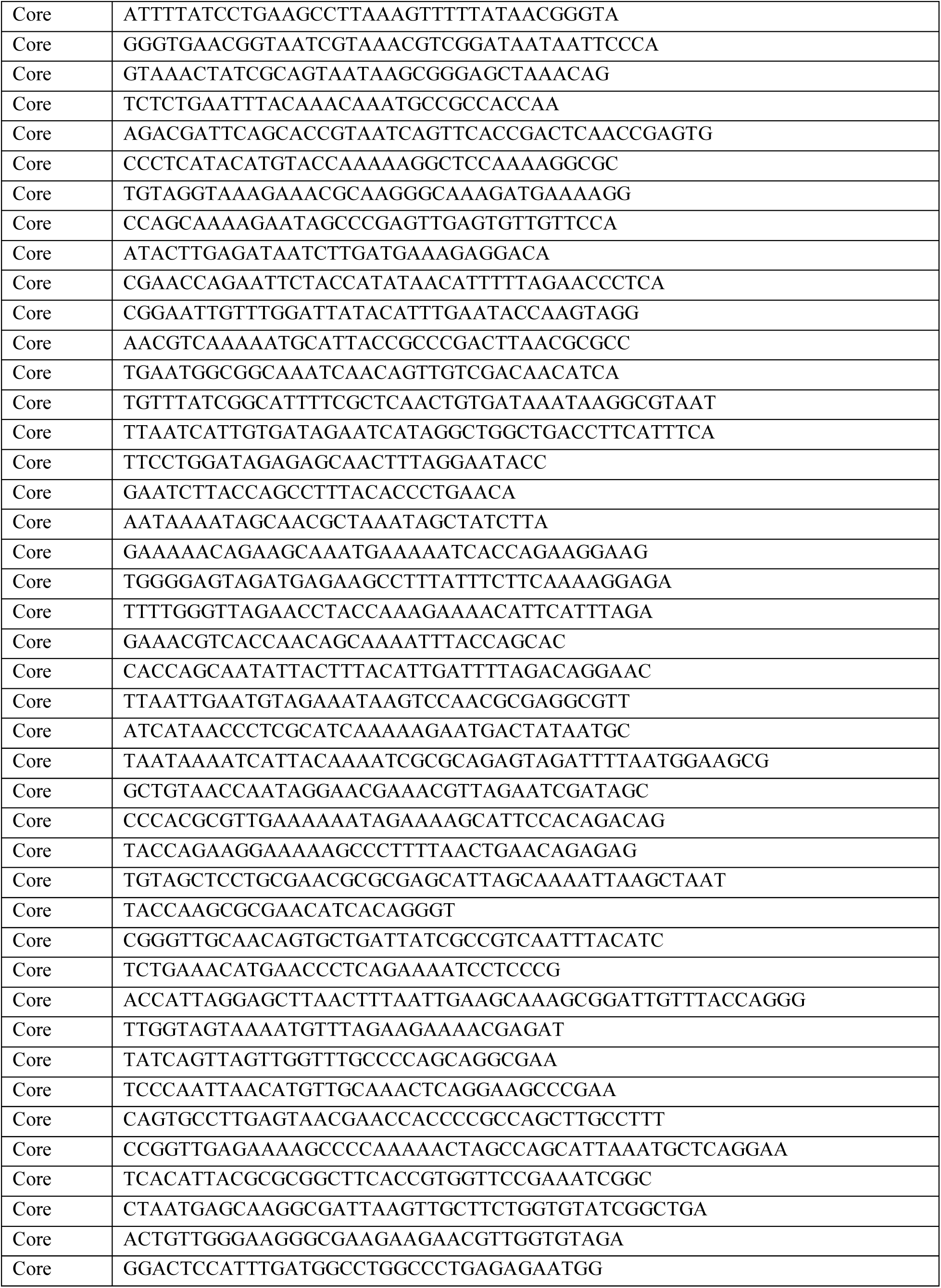
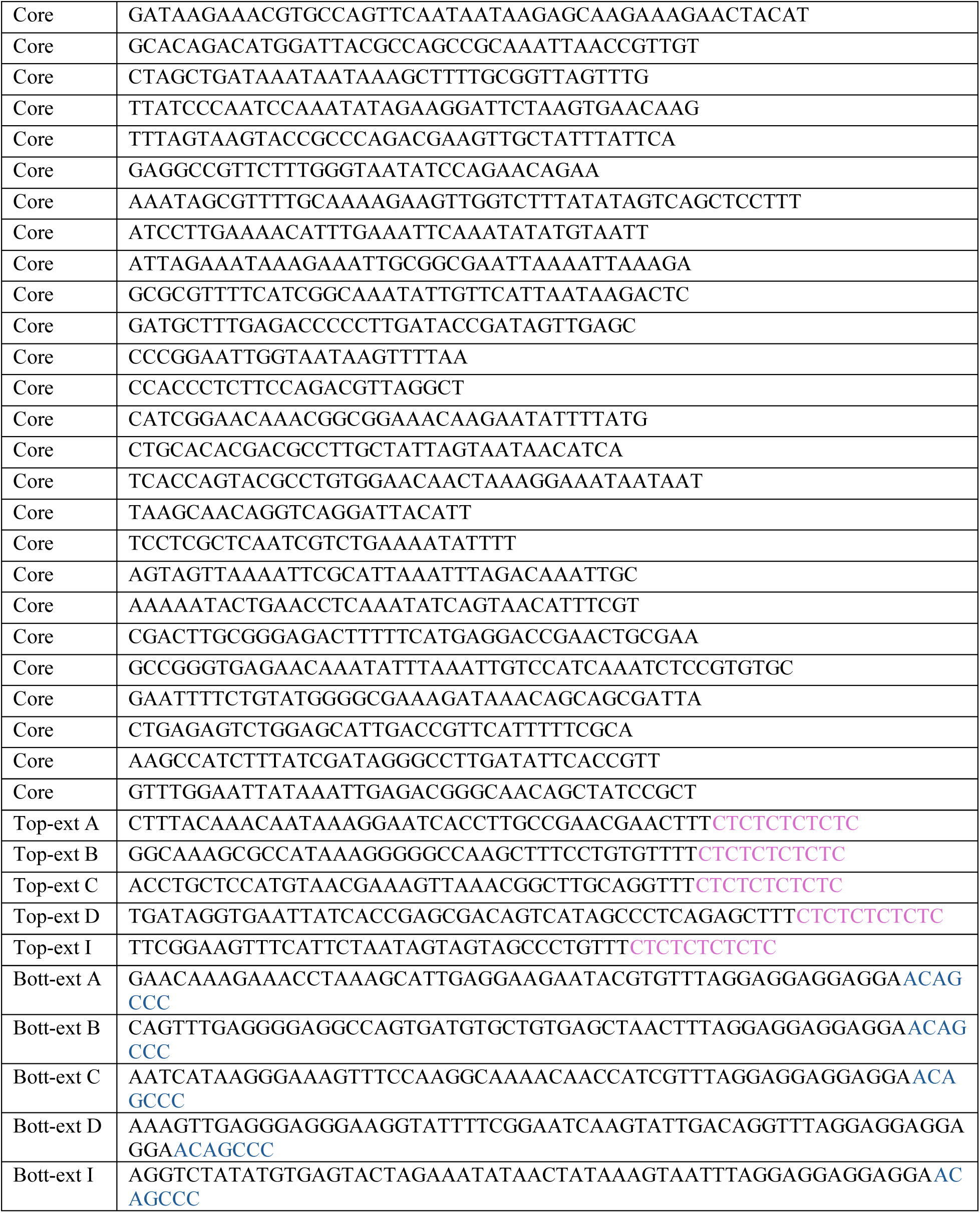
Sequences of core staples of the origami stamp.

**Table S6.**
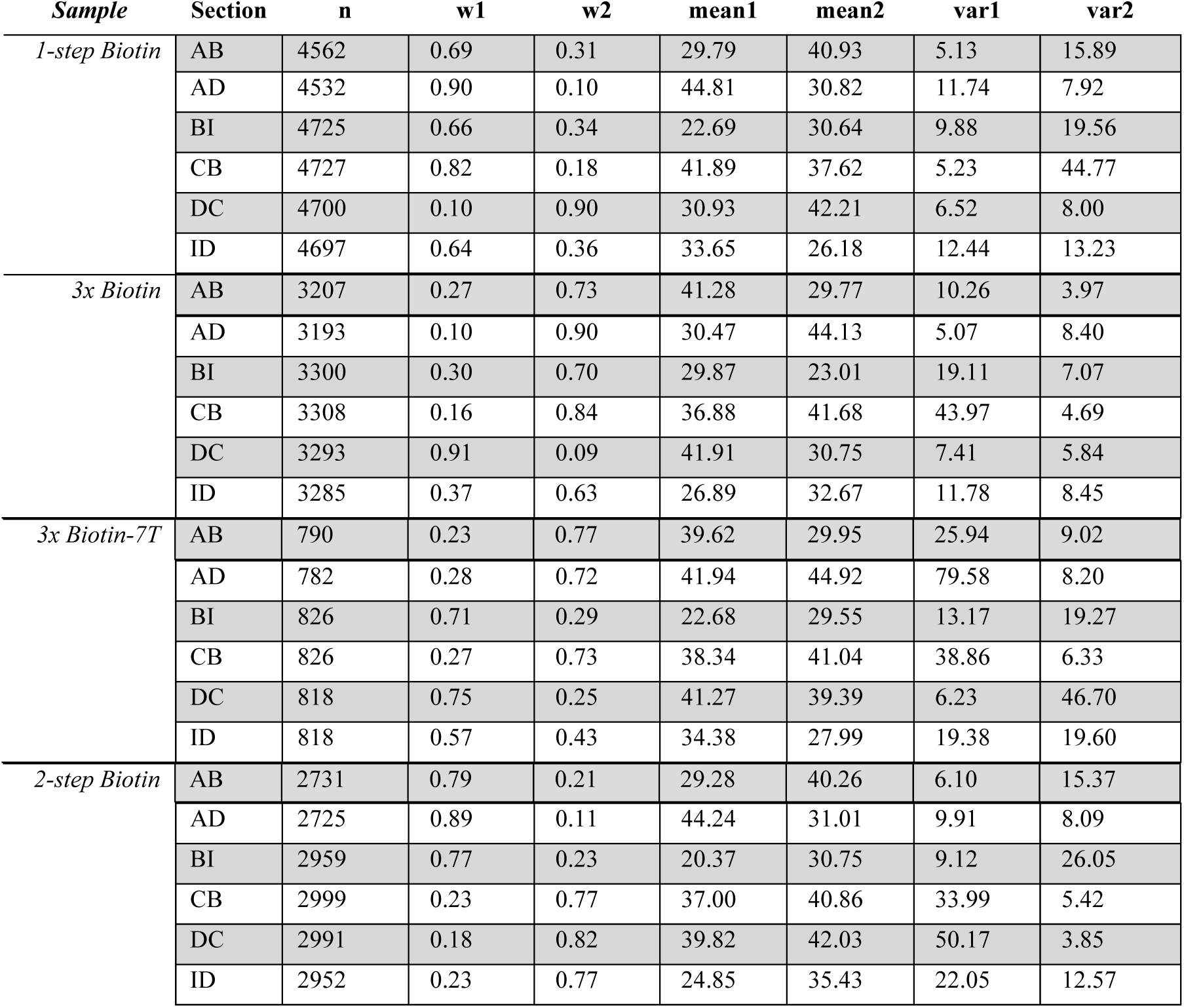
Parameters derived from the Gaussian mixture model decomposition. The segments were measured across the different stamps and the distributions were decomposed using a two-component Gaussian mixture model. The number of input observations (n) yielded the weights (w1 and w2) as well as means (mean1, mean2) and variances (var1, var2) for each Gaussian component respectively.

## Notes

### Competing Interest Statement

The authors have declared no competing interest.

