## Supplemental information for "Nanoscale precise stamping of biomolecule patterns using DNA origami"

### Table of Contents

Supplemental Figures 3

Figure S1. TEM images of the origami stamp. 3

Figure S2. Toehold-mediated strand displacement mechanism. 4

Figure S3. SPR-monitored stamping mechanism. 5

Figure S4. Signal of non-filtered origami stamps and single-stranded PTOs in magnified FOV. 6

Figure S5. Conjugation of 3xBiotin-PTO and 3xBiotin-7T-PTO. 7

Figure S6. View of 3xBiotin and 3xBiotin-7T stamping. 8

Figure S7. Data Processing. 9

Figure S8. Gaussian mixture model decomposition of the distributions of measured distances after rotational alignment. 10

Figure S9. Filtered distributions of the measured distances. 11

Supplemental Tables 11

Table S1. Averaged localization precision (nm) calculated by Nearest Neighbor Analysis (NeNA) 11

Table S2. Percentage of fully, partially, and failed stamped patterns 11

Table S3: Summary statistics of the different segments and the deviation to the designed distances. 13

Table S4. Sequences of staple extensions, PTOs, invader, and imager strands. 13

Table S5. Sequences of core staples of the origami stamp. 14

Table S6. Parameters derived from the Gaussian mixture model decomposition. 20

### Supplemental Figures

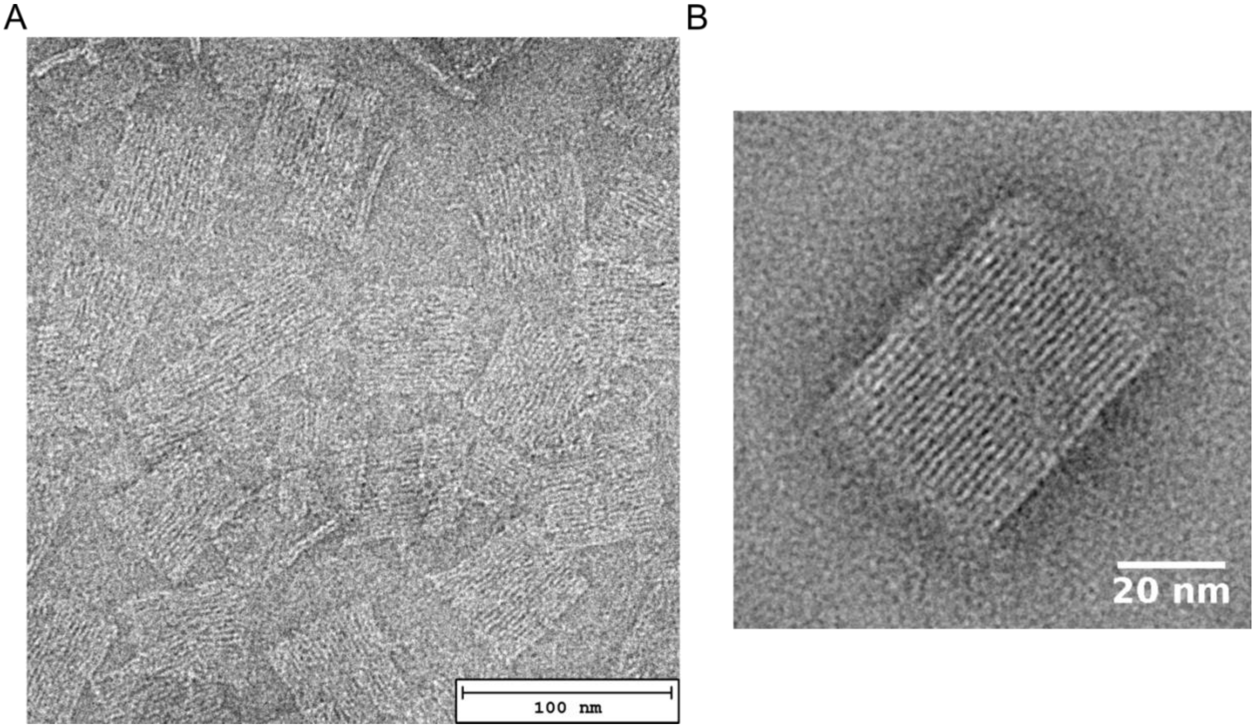


**Figure S1. TEM images of the origami stamp.** (A) Raw image of origami stamps on a grid. (B) Scipion software 2D rendering of the origami stamp upon averaging of over 100 manually selected structures.

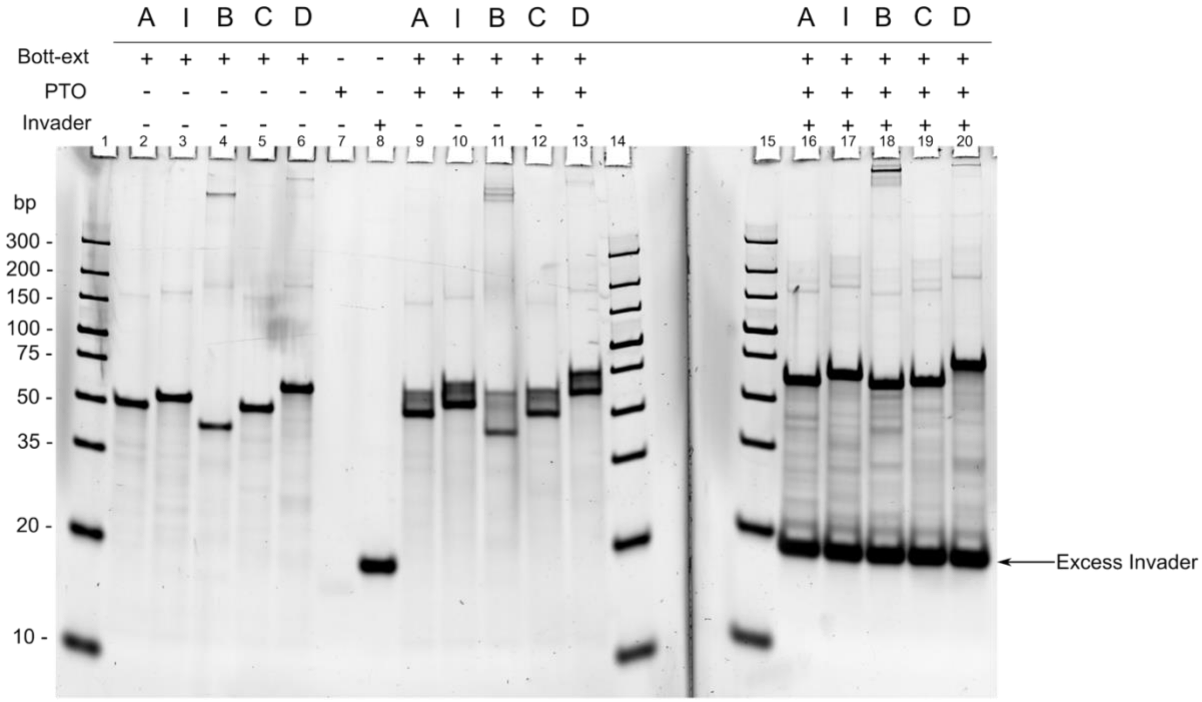


**Figure S2. Toehold-mediated strand displacement mechanism.** 10% acrylamide native gel showing Bott-ext oligonucleotides (wells No. 2-6), biotinylated PTO (well No. 7), and the invader sequence (well No. 8). PTO were annealed to each single Bott-exts and the complexes were visualized in the same gel (wells No. 9-13) as an upward shift. Part of the Bott-ext-PTO complexes were mixed with excess invader sequence and loaded in the gel (wells No. 16-20). A larger upward shift was observed in this case, given the longer length of the invader oligonucleotide, which displaced the PTO and annealed to the Bott-exts from their 3' toehold. The gel was stained with SYBR gold stain. Unfortunately, the single PTO and the strand-displaced PTOs (wells No. 16-20) were poorly visible or invisible, as the PTO is not efficiently stained by SYBR Gold.

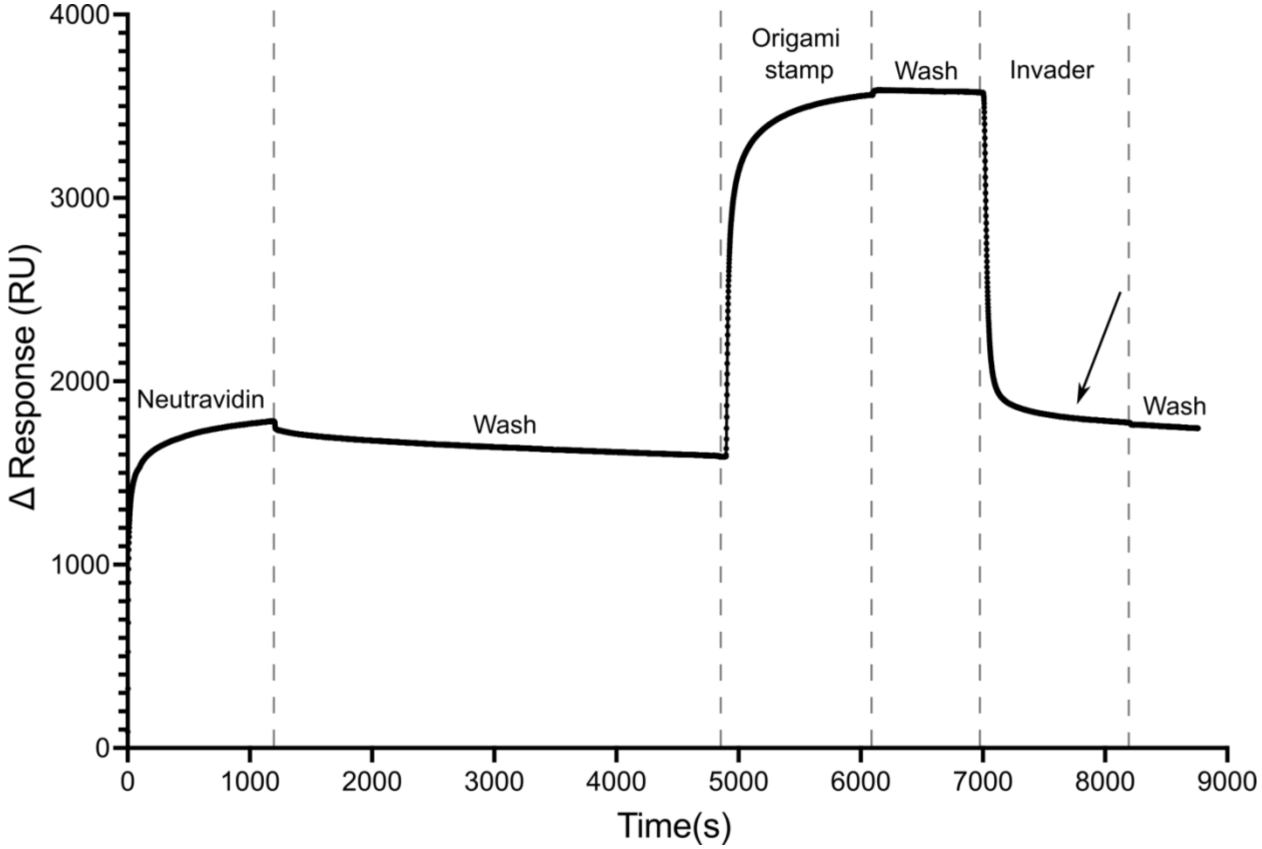


**Figure S3. SPR-monitored stamping mechanism.** Molecular stamping mechanism characterization by SPR, where the PEG-b-neu surface was generated by coating the PEG-biotin brush with neutravidin (from 0 to 5000 s), followed by the immobilization of the origami (from 5000 to 7000 s), indicated by an increase in the response. The application of the invader oligonucleotide (from 7000 s) caused a quick decrease in the response, indicating the release of the origami structures in solution. The final response (black arrow) indicated the presence of residual signal after the invasion, suggesting the presence of bound stamped PTOs, as a result of the complete stamping process.

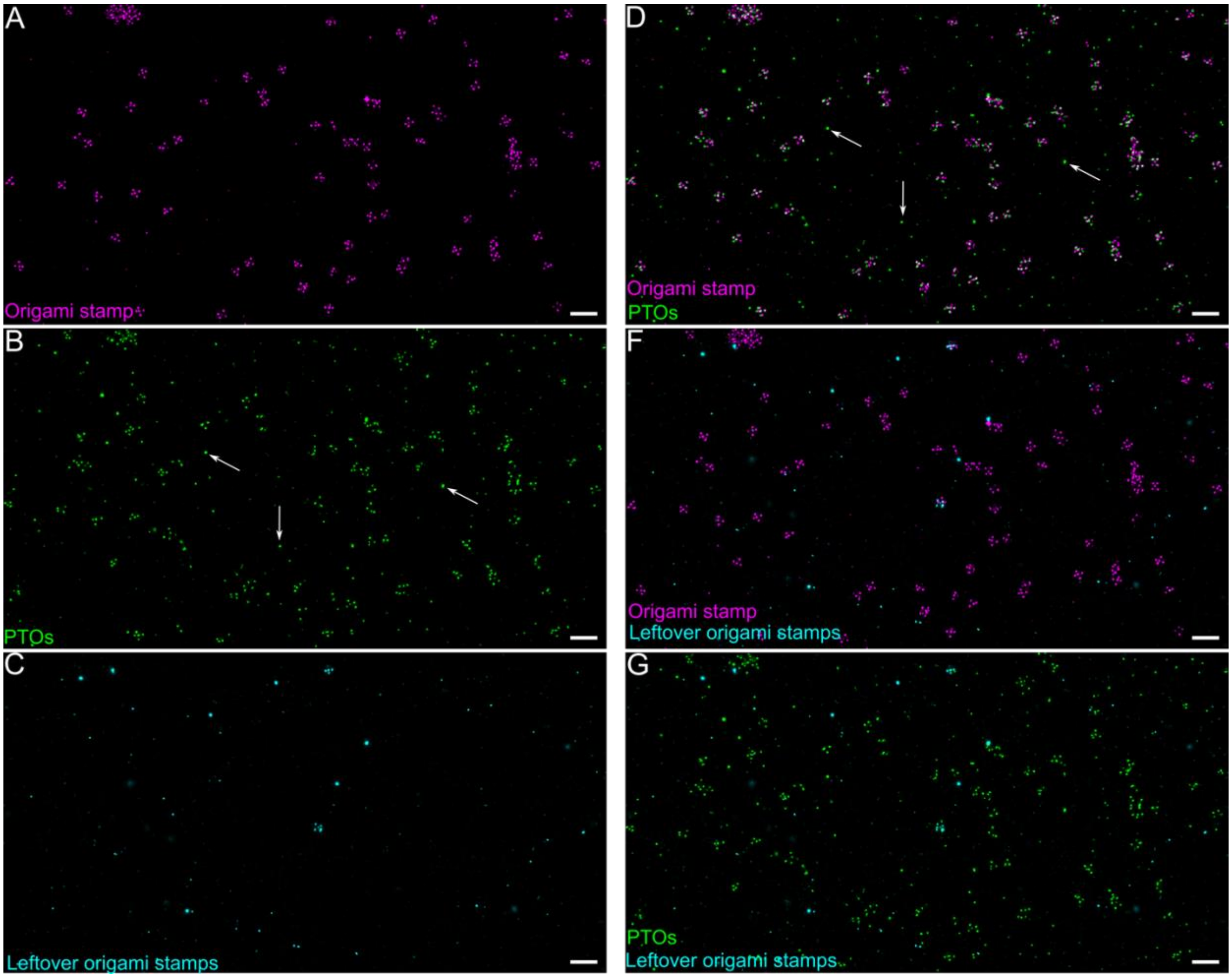


**Figure S4. Signal of non-filtered origami stamps and single-stranded PTOs in magnified FOV.** (A) Non-filtered signal of origami stamps; (B) Non-filtered signal of single-stranded PTOs. White arrows indicate PTO signals unrelated to stamped patterns; (C) Non-filtered signal of origami stamps, visualized after the stamping mechanism, showing the leftover origami stamps not successfully released in solution during strand displacement; (D) Merged signals of origami stamps and single-stranded PTOs. White arrows indicate PTO signals unrelated to stamped patterns; (E) Merged signals of origami stamps and leftover Origami, after stamping, and (G) Merged signals of single-stranded PTOs and leftover origami. Scale bar 200 nm.

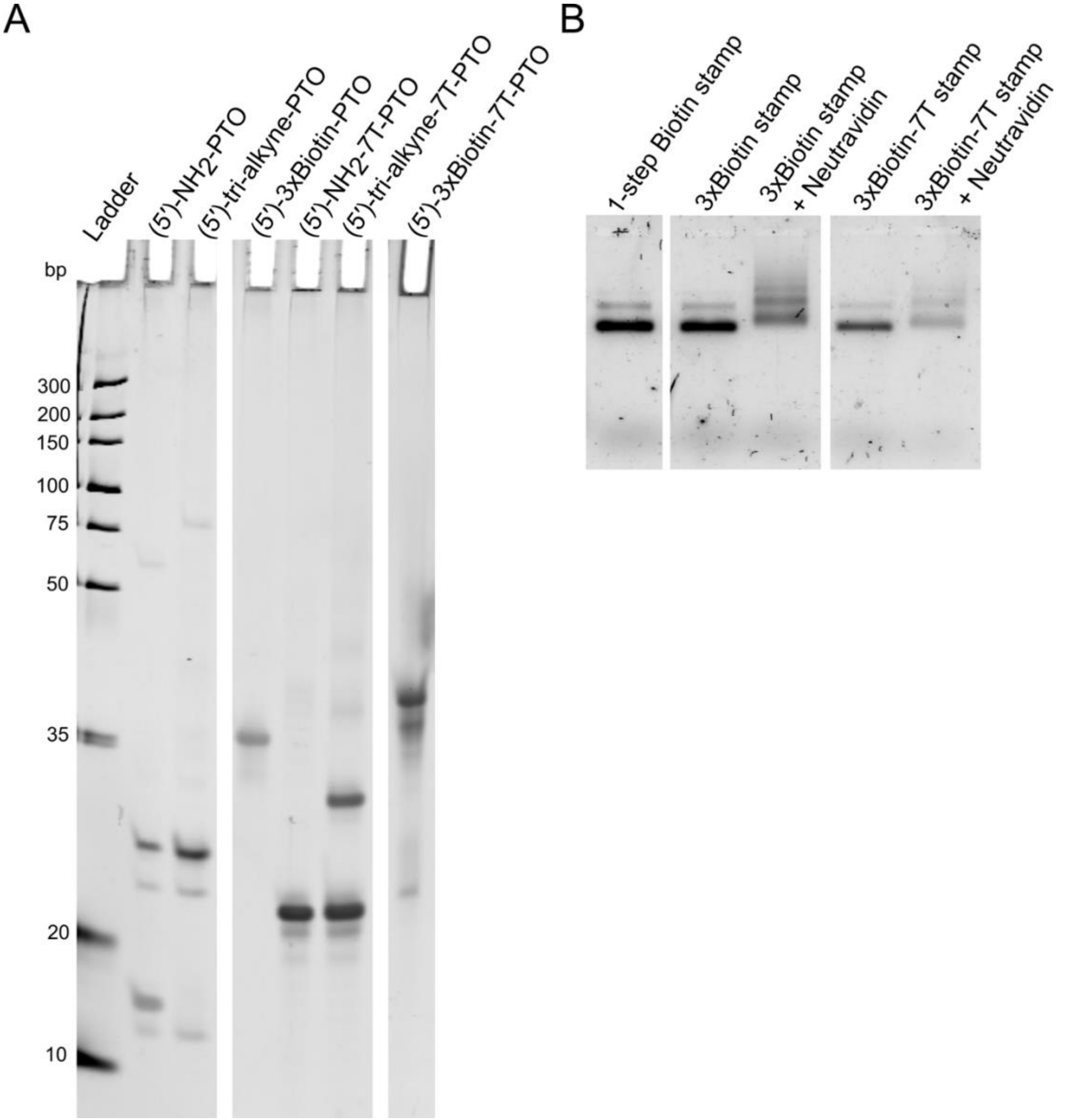


**Figure S5. Conjugation of 3xBiotin-PTO and 3xBiotin-7T-PTO.** (A) A 16% denaturing PAGE gel of 5′-$NH_2$-PTO and 5′-$NH_2$-7T-PTO conjugated to tri-alkyne linker and successively functionalized with three biotin molecules. (B) origami stamp assembled either with biotin-PTOs (1-step Biotin stamp), 3xBiotin-PTOs (3xBiotin stamp) and 3xBiotin-7T-PTOs (3xBiotin-7T stamp). The last two origami were incubated with excess neutravidin to verify by mobility shift assay the presence of biotinylated PTO annealed to the stamp.

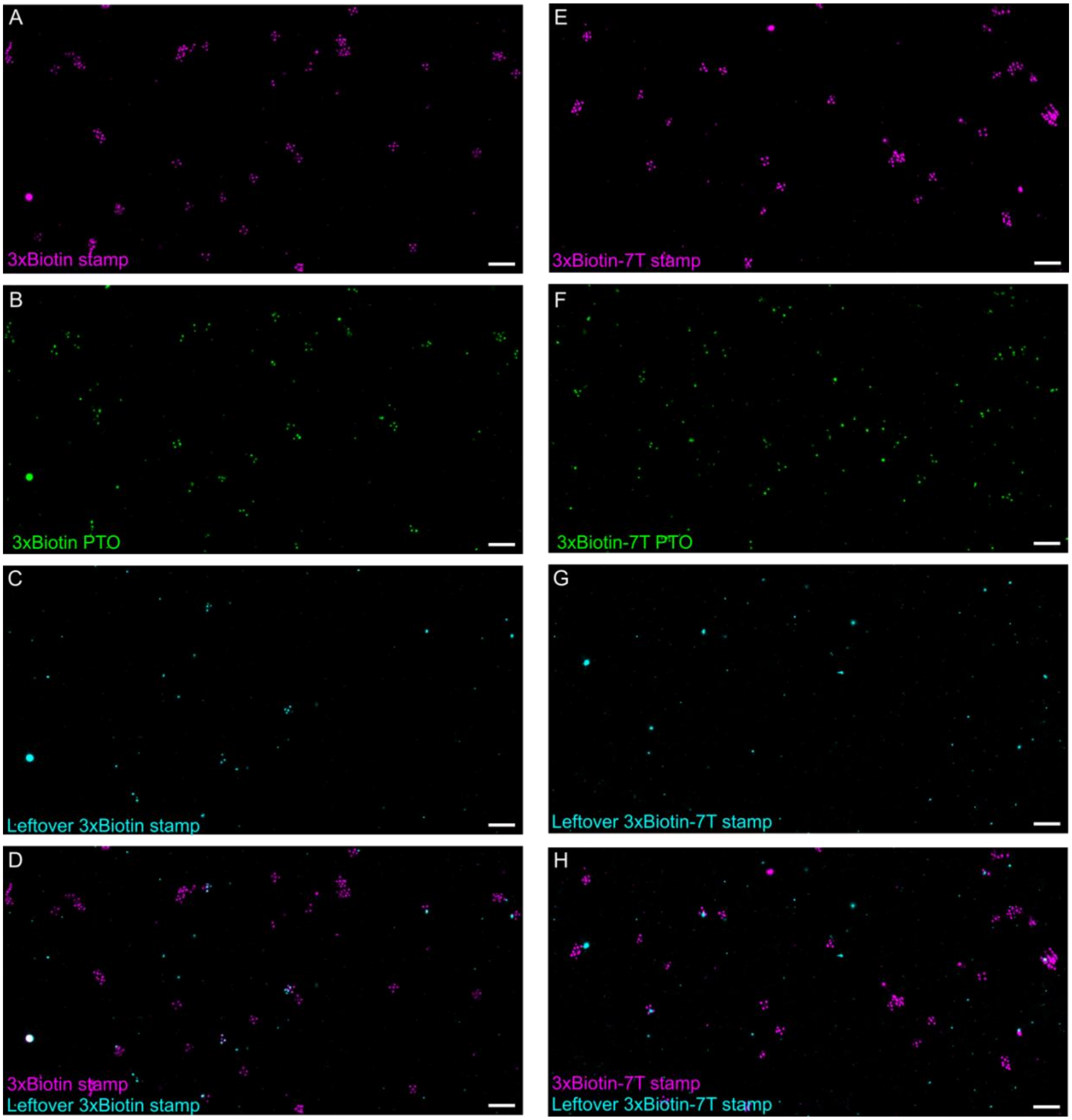


**Figure S6. View of 3xBiotin and 3xBiotin-7T stamping.** (A) Non-filtered signal of 3xBiotin stamps; (B) Non-filtered signal of 3xBiotin PTOs; (C) Non-filtered signal of leftover 3xBiotin stamps; (D) Merged signal of 3xBiotin stamps and leftover 3xBiotin stamps after strand displacement; (E) Non-filtered signal of 3xBiotin-7T stamps; (F) Non-filtered signal of 3xBiotin-7T PTOs; (F) Non-filtered signal of leftover 3xBiotin-7T stamps; (G) Merged signal of 3xBiotin-7T stamps and leftover 3xBiotin-7T stamps after strand displacement. Scale bar 200 nm.

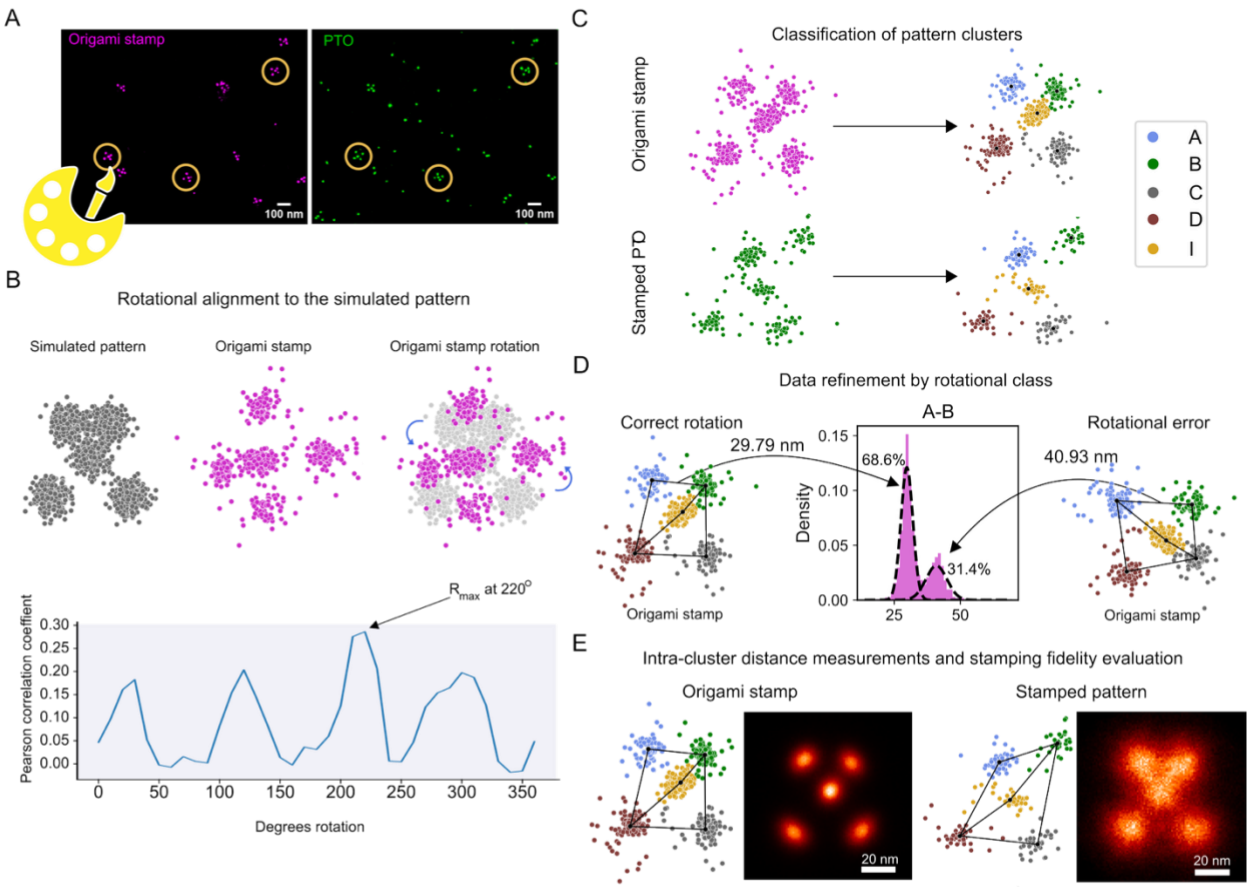


**Figure S7. Data Processing.** (A) Origami patterns were identified and selected in Picasso Render, and the corresponding localizations of the origami stamp and PTOs were exported for processing in the custom script. (B) The origami stamp localizations were rotated and aligned onto a simulated pattern. The angle of rotation was determined from the maximum Pearson correlation coefficient (bottom). (C) The aligned localizations were further classified using a simulated distance reference to assign the most probable strand of origin for each smaller cluster in each ROI. The cluster centroid is marked in black. (D) To identify rotational errors, the center of mass for each Top-ext cluster was used as the point of origin for the determination of intra-cluster distances. Each distance was analyzed using a two-component Gaussian mixture model (Figure S8) to obtain rotational models and identify the set of erroneously rotated patterns. Patterns containing rotational errors were filtered out, and the remaining origami stamp-stamped pattern pairs were used in the final ensemble. (E) Intra-cluster distances were measured for all the Top-exts and PTOs pattern pairs to determine the stamping fidelity of each origami stamp.

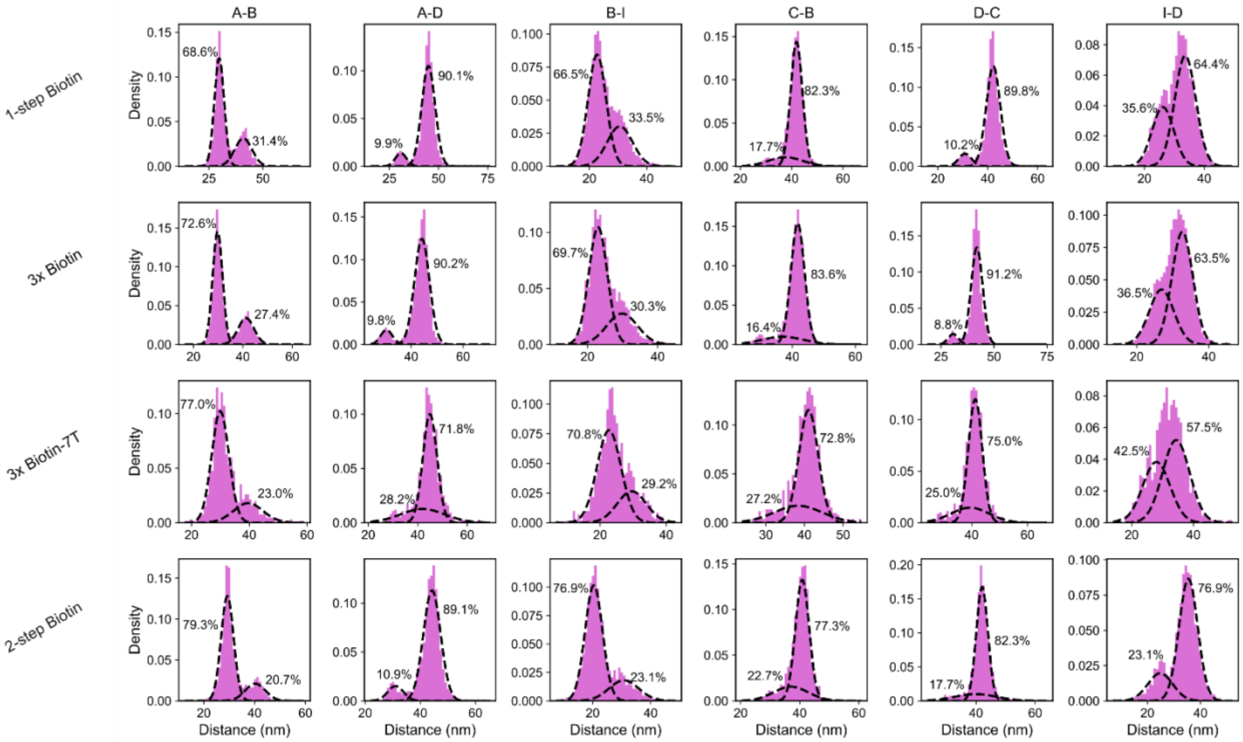


**Figure S8. Gaussian mixture model decomposition of the distributions of measured distances after rotational alignment.** Across each origami stamp, the measured distances between the observed Top-exts were analyzed using a two-component Gaussian mixture model. The weights are shown as percentages next to the respective component, and the parameters of the derived components are provided in Table S6. The resulting models were used to distinguish and filter erroneously rotated origami stamps and their corresponding PTO from the ensemble, as shown in Figure S7.

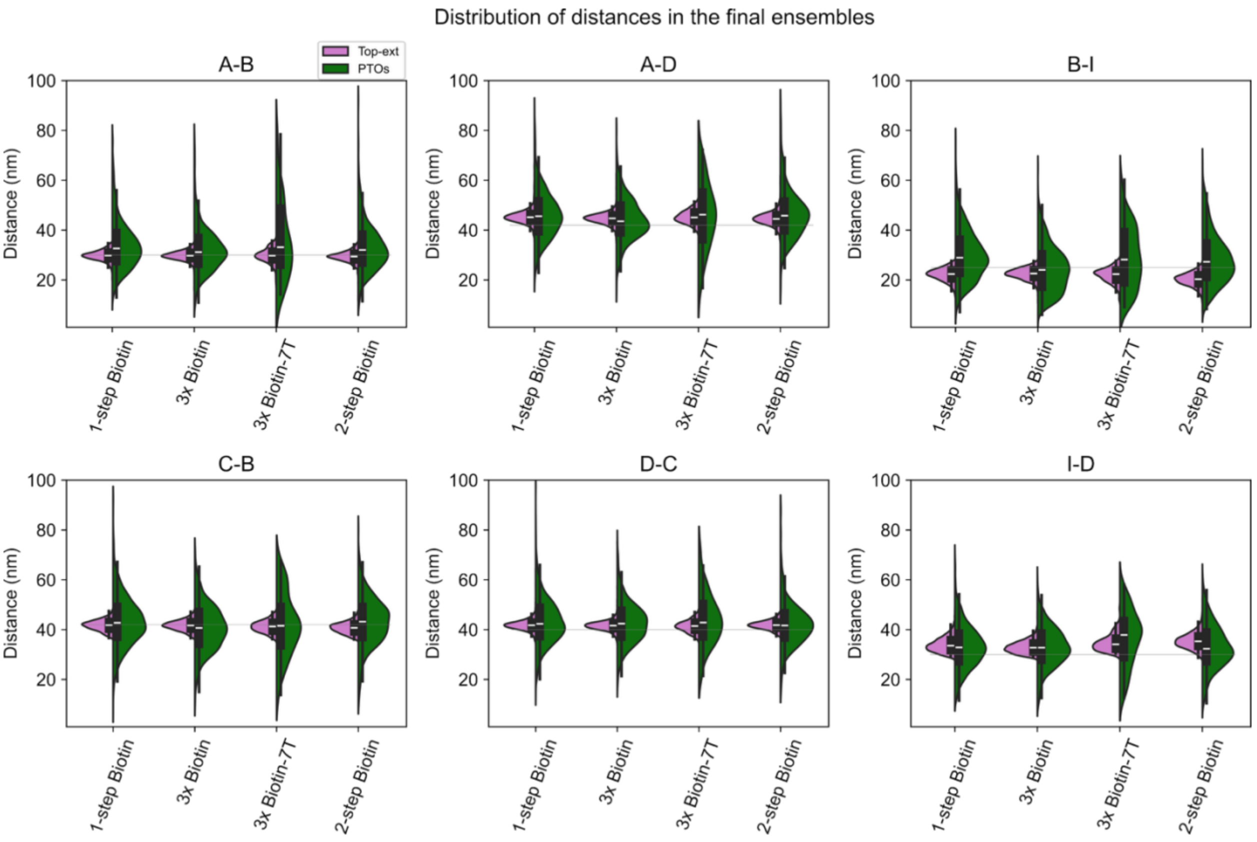


**Figure S9. Filtered distributions of the measured distances.** Violin plots and boxplots of the measured distances between Top-exts of the origami and PTOs after data filtration according to the rotational model in Figure S8. The white line and boxes show the median and IQR. The horizontal grey lines represent the designed distances.

### Supplemental Tables

**Table S1. Averaged localization precision (nm) calculated by Nearest Neighbor Analysis (NeNA)**

| *Origami stamp* | *Imaged Top-exts (nm)* | *Imaged Stamped PTOs (nm)* |
|---|---|---|
| 1-Step Biotin | 2.98 | 2.94 |
| 3x Biotin | 4.34 | 3.81 |
| 3x Biotin-7T | 4.22 | 3.52 |
| 2-step Biotin | 3.81 | 3.60 |

**Table S2. Percentage of fully, partially, and failed stamped patterns**

| *Origami stamp* | *Stamped PTO* | *Percentage (%)* | *Number of observed patterns* |
|---|---|---|---|
| *1-step Biotin stamp* | 0 | 2.05 | 52 |

| | | | |
|---|---|---|---|
| $n_{tot}$ =2536 | 1 | 1.77 | 45 |
| | 2 | 12.5 | 317 |
| | 3 | 33.6 | 852 |
| | 4 | 36.2 | 918 |
| | 5 | 13.88 | 352 |
| *3x Biotin stamp* $n_{tot}$ =1886 | 0 | 0.21 | 4 |
| | 1 | 2.7 | 51 |
| | 2 | 15.91 | 300 |
| | 3 | 34.62 | 653 |
| | 4 | 33.67 | 635 |
| | 5 | 12.88 | 243 |
| *3x Biotin-7T stamp* $n_{tot}$ =348 | 0 | 6.32 | 22 |
| | 1 | 17.53 | 61 |
| | 2 | 35.06 | 122 |
| | 3 | 25.29 | 88 |
| | 4 | 12.36 | 43 |
| | 5 | 3.45 | 12 |
| *2-step Biotin stamp* $n_{tot}$ =1856 | 0 | 1.19 | 22 |
| | 1 | 4.74 | 88 |
| | 2 | 25.22 | 468 |
| | 3 | 37.66 | 699 |
| | 4 | 25.38 | 471 |
| | 5 | 5.82 | 108 |

**Table S3: Summary statistics of the different segments and the deviation to the designed distances.**

The segments were measured across the different stamps and their stamped PTOs. The median, mean and standard deviations (stdev) values are shown to provide a reference of the physical scale of the stamping. The designed distances are used as references to obtain the absolute average distance deviation from the experimental mean values. All values are in nm.

| *Origami stamp* | *Segment* | *Top-ext median* | *Top-ext mean* | *Top-ext stdev* | *PTO median* | *PTO mean* | *PTO stdev* | *Designed distance* | *Top-ext deviation* | *PTO deviation* |
|---|---|---|---|---|---|---|---|---|---|---|
| *1-step Biotin* | AB | 29.8 | 29.8 | 2.0 | 32.7 | 34.1 | 10.0 | 30.0 | 0.2 | 4.1 |
| *3x Biotin* | AB | 29.8 | 29.8 | 1.8 | 31.2 | 32.2 | 9.1 | 30.0 | 0.2 | 2.2 |
| *3x Biotin-7T* | AB | 29.8 | 29.8 | 2.8 | 33.1 | 36.1 | 15.5 | 30.0 | 0.2 | 6.1 |
| *2-step Biotin* | AB | 29.4 | 29.4 | 2.1 | 32.1 | 33.2 | 9.7 | 30.0 | 0.6 | 3.2 |
| *1-step Biotin* | AD | 45.2 | 45.3 | 2.6 | 45.6 | 46.1 | 9.6 | 42.0 | 3.3 | 4.1 |
| *3x Biotin* | AD | 44.7 | 44.7 | 2.2 | 43.5 | 44.3 | 8.5 | 42.0 | 2.7 | 2.3 |
| *3x Biotin-7T* | AD | 45.1 | 45.3 | 2.5 | 46.2 | 45.8 | 13.3 | 42.0 | 3.3 | 3.8 |
| *2-step Biotin* | AD | 44.5 | 44.5 | 2.5 | 45.8 | 46.4 | 9.8 | 42.0 | 2.5 | 4.4 |
| *1-step Biotin* | BI | 22.4 | 22.1 | 2.6 | 28.9 | 30.0 | 9.9 | 25.0 | 2.9 | 5.0 |
| *3x Biotin* | BI | 22.6 | 22.5 | 2.2 | 24.0 | 24.4 | 9.3 | 25.0 | 2.5 | 0.6 |
| *3x Biotin-7T* | BI | 22.3 | 21.9 | 3.0 | 28.2 | 29.2 | 12.3 | 25.0 | 3.1 | 4.2 |
| *2-step Biotin* | BI | 20.3 | 20.2 | 2.6 | 27.3 | 28.7 | 10.3 | 25.0 | 4.8 | 3.7 |
| *1-step Biotin* | CB | 41.8 | 41.7 | 2.1 | 42.8 | 43.3 | 10.0 | 42.0 | 0.3 | 1.3 |
| *3x Biotin* | CB | 41.7 | 41.7 | 2.1 | 40.7 | 40.8 | 9.2 | 42.0 | 0.3 | 1.2 |
| *3x Biotin-7T* | CB | 41.3 | 41.3 | 2.3 | 41.5 | 42.1 | 12.5 | 42.0 | 0.7 | 0.1 |
| *2-step Biotin* | CB | 40.7 | 40.7 | 2.2 | 43.2 | 43.2 | 9.7 | 42.0 | 1.3 | 1.2 |
| *1-step Biotin* | DC | 41.8 | 41.8 | 2.1 | 42.3 | 43.2 | 9.1 | 40.0 | 1.8 | 3.2 |
| *3x Biotin* | DC | 41.6 | 41.6 | 2.2 | 42.4 | 42.7 | 8.2 | 40.0 | 1.6 | 2.7 |
| *3x Biotin-7T* | DC | 41.4 | 41.4 | 2.3 | 42.8 | 43.9 | 10.6 | 40.0 | 1.4 | 3.9 |
| *2-step Biotin* | DC | 41.8 | 41.8 | 1.8 | 41.7 | 42.1 | 8.8 | 40.0 | 1.8 | 2.1 |
| *1-step Biotin* | ID | 33.6 | 34.0 | 3.0 | 32.9 | 33.2 | 8.7 | 30.0 | 4.0 | 3.2 |
| *3x Biotin* | ID | 32.8 | 33.0 | 2.4 | 32.8 | 33.0 | 8.0 | 30.0 | 3.0 | 3.0 |
| *3x Biotin-7T* | ID | 34.1 | 34.7 | 3.2 | 37.9 | 36.3 | 11.3 | 30.0 | 4.7 | 6.3 |
| *2-step Biotin* | ID | 35.3 | 35.5 | 3.1 | 32.3 | 33.2 | 8.9 | 30.0 | 5.5 | 3.2 |

**Table S4. Sequences of staple extensions, PTOs, invader, and imager strands.**

| **Module** | **DNA sequence (5′-3′)** |
|---|---|
| Origami Top-ext (3xR3d) | Top staple-TTTCTCTCTCTCTC |
| Origami Bott-ext (toehold sequence) | Bottom staple-TTTAGGAGGAGGAGGAACAGCCC |
| PTO (3xR1d) | biotin-TTTTCCTCCTCCTCCT |
| 7T-PTO (3xR1d) | TTTTTTTTTTTCCTCCTCCTCCT |
| Invader | GGGCTGTTCCTCCTCCTCCTAAA |
| R3i-Cy3B | GAGAGAG-Cy3B |

| | |
|---|---|
| R1i-Cy3B | AGGAGGA-Cy3B |

**Table S5. Sequences of core staples of the origami stamp.**

| Module | DNA sequence (5′-3′) |
|---|---|
| Core | GAATATAGTTCAGCTTCCAAGTTTCATCGTAGG |
| Core | CCACCACCCCCTTATTAGCGTTTGGGCGACATTTGA |
| Core | TAGGCAGAAACAATAGACCAATCAAGCAAGCAAATCA |
| Core | GAGGCCACCGAGTAAAAGAGTCTGAGAA |
| Core | TCGTAATCGTTGCAGCCAGCTGCAAGGGCGAAAAACCGTC |
| Core | TCGGGCTGGCCACCGAGCTCGAAT |
| Core | AATCACCGGAACCTCATATGGTCACCAGTAGT |
| Core | TTAGCCTAATTATTTTTGCGCTAATATCAGAGAGAAATACATGGCA |
| Core | GTACTTGCTGAACATAAATCAAAAATCATTGCCAGAACGACGATTTA |
| Core | GCGAGTAACAACCACTAGCATTGTATAAGAGGCCGGA |
| Core | TTAAGGAAAGCGGCCACCCTCAGAGCCA |
| Core | AATTTTAAAAGTTTGAAACCCTCAGGTCAGTTTATTAGTC |
| Core | GGGTAAGAAGAAAAATCTACGTACGAGAAAAATCAACGGTC |
| Core | AGCTGAAACCTCATAATCCTCAGAACCGCCACTATT |
| Core | GCATTAGATTTGTCACCCGAACAAAGCACCATTACACAAA |
| Core | TGTCGTGCAAGCGGTCCACGCAGAAGAACTCATTAACACCGCCA |
| Core | GAAATTGTTGATTGCCGGAGAGGCCACTATTAAAGAACGT |
| Core | TCGGGTAAGCGTCGCCACCCTCAGAACCGCCACCCAGGATTAGTAT |
| Core | TATTAACGTCAGATGAATACTTTGAATTACCTTGTA |
| Core | TCTGAGAGACTACAAATCCGAATAAAAATTCTT |
| Core | AAAATCCCCAAGAGTCGGTTTGCGTATTGGGCTGGGGTGC |
| Core | TTAAACCATCATATGCACAAAAGGTATGTAAATGCTG |
| Core | GACAGTCATGTAATACCTCAGAGCTGTTTAGCTATATTTTGAGA |
| Core | TATTTTTGAGAGCAGGCAAGATAAAAAGTTGAT |
| Core | AGGTTGATGATACAGTATAAACAATAAGTGCCGTCGAGAGGGTTGAT |
| Core | TCCCACGCTGACACTAACAAGATAGAACCCTT |
| Core | AGCAATACATTAAAGGGGCAGATTCACCAGTACCT |
| Core | AAGGAAACGCAAACGCAATTTTGGGAATTAG |
| Core | CGCCACCAAGTGCCCGGAGTGTACAGGTGTATCACCGTAC |
| Core | TTGTACAGTAATAATCCTGATTATCATCATAT |
| Core | GGGTTGTTAAATCAGCAATGGGATAGGTCTGATGGCAATTACAT |
| Core | CTTGCCTGAGAATCAGAAAGGGACATTAAACC |
| Core | ATCCTAATTTACGAGCGAATCGCCACGCCAACATTTTAGT |
| Core | TGTCGTCTATTTTCAGTCGGTTTATCAGCTTGCTTTCAATGAGTAAAT |
| Core | AGTATTTAGTACCATACATGGCTTTTGAGGCAGGTC |
| Core | TAGGAACCGTTAGCGTTTTCAACAGTTTCAGCGGCCGCTTAATG |
| Core | TGTTAGCAAACGTAGAATAACCCAGTTAAGCCACAAAATA |
| Core | TATATTTAAAGATTAACATCCAATAAATCATAATCTACAACCTG |

| | |
|---|---|
| Core | CCTTTTACATTTAACAATTTCTTCTGAATCAGGTTTAGA |
| Core | ACTAATTTTAAGTACTGCGGAATCGTCATAAATTGCGGAACGCC |
| Core | AGAGTCAATAGTGAAACTTTTACCGACCGAGTAGGGC |
| Core | TTGAATCCATCGCGTTTTAATCATCGAGAACAAAAAACAGACCC |
| Core | GGAATAAGTTTATCGGGAGAATTTAAGAAACA |
| Core | GCAACATATAAAATCAGAGGGAAATAGCACGA |
| Core | TGGCATCACCGGAATTAAATATGCTTTAAACAG |
| Core | TTTTTCACATAACCGACGCTGAGGGTAAAATACGTAATGCGAAGGCAC |
| Core | GTACAACGTACAGACCAGGCGCAA |
| Core | GACCTAAATTTAATGGTAGCGATAAGACGCTGCAATTACC |
| Core | CACAATTCGACGTTGTAAAACGACCGACGACAGCCG |
| Core | GCGTTAGCTGTTGCATGCCTGCAGGTCTCGTAACCAG |
| Core | TTACAGGTAGAAAATTACCCACACCAGAAGCG |
| Core | AGCTAAGTAAGCAGAATTACCTTATGCGATGCAGATACATAAGC |
| Core | TGATAGCCCTAAAACACGCTCATGCATTTTGAGAGAAGTGTTTTTATA |
| Core | TGATAAGATTTTGCGGATGGCTTAATACATTTCAATAACCATAAAGCT |
| Core | TGATGAAACAAACATCTATCAAAAGTAAAACAAATCCTTT |
| Core | ACAAGAATACACTAAAACACAATTGTGTACCA |
| Core | ACTTCAAGAACCGGATATTCGATTCATCGACG |
| Core | CCAACGAGCCAGGACAAAGAACGCGAGAAATTTATCTTTC |
| Core | TCAGGAGGCCAGGCGGGTTAATGCCCCCTGCCCCTCAGAG |
| Core | ATCCCGAGTAGTAAATTGGGCCAGTCAGAGTTGAGAACT |
| Core | TCATGCGCTCTTACGCCAGCTGGCGTCGCCATTCGTG |
| Core | CTTTTAAACAACAACGATCTAAAGTTT |
| Core | GAATTTCTCAGCATCGGAACGCGGAACCGCCTAGTA |
| Core | AGCAACGGCTACAGAGGAACGGTGGAGATTTGTTTCAACT |
| Core | AGACATAGTACGTCCAAAACTGGCTCATT |
| Core | ATACGAGCCATAAAGTGTAAAGCCGCCAGGGTTTCA |
| Core | AATAACATAGCAAGCCTCAAGATTCGACAATAAACAA |
| Core | TGGGCGCAGACTCTAGCTTCGCTAACTGCCCGCTTTCCAG |
| Core | ATGCCTTTTTATTGCTTCTTTTTAATGGAAAC |
| Core | GATCGCACCAGCTTTCCGGCACCGGGTAACGCCCAGTCACCACACAAC |
| Core | CCAAACCTATGATTAGCGGGGTTTTGCTC |
| Core | ATTGAATAACCACCTCCGGCGGATTCGCCTGA |
| Core | GAAAGCGTCTGGCCTTCCTGAGGAAGATGTCAATCAGTT |
| Core | TAGAAAATAGAGCCACTAGCAAGGATTAAAGCCAGGAGGT |
| Core | TGCAATTGTAGGATAGCAAGCCCAA |
| Core | GAGTTAAAGGAGTGAGTCTCCAAAGTAACACTGAGTTTCG |
| Core | CAACCTAATACTTAGCCGCAGACGTAACAAAGCTGCTCAT |
| Core | ACCAGTATCCTTATCATAATGCAGGCGGGAGGTTTT |
| Core | AAAAGGAATTACGAGGCTTCAAATCCCTCAAATGCAACTA |
| Core | ATATCATCTTTGGACTAAATCGTCACCCTCAGCAATTT |
| Core | GCGTCTTTCCAGAGCGAACCTCGCCCAATATAATCGGCTGTCTTTAAAG |

| | |
|---|---|
| Core | AAATAACAATAACTTAGGTTGGGTTTAAATGCAATGAGGCTATC |
| Core | AGTCATCAATACAGTACCTTAGATAATACATT |
| Core | CATAAGTACCGGTTATACACACCGGAATCATA |
| Core | TGATTTAGGAGGAGCCAGCGGTGAGGCGGTCA |
| Core | CCGCCGAGGAAAGACACCGCCAAAGACAA |
| Core | ACAGACTAATAGATTTCGGTGCGGGCCTAGGATCCC |
| Core | AATCCTGACGTCAATTAATGAATCGGCCAAATT |
| Core | GGTACGCCTCCATCACATTGCAACAGGAAAAATCGCCATT |
| Core | AAGTACGGTGTCTTGTACTTCGAGCTTCAAAG |
| Core | AATAAGGCTTGCCCTGTAATAAAAGAACAACAAAAAACCA |
| Core | GCCAAATAACGGAATACCCAAAACAATGTAATTGAGTTT |
| Core | GAGACTCCTCAAGAGATCAGAACCCAG |
| Core | AATCGTCGCTATTTAAATAAAATCGCAATAATAAGA |
| Core | AAATCGGTAAAAACATTATGACCCAATCACCAATTCAACCTATGTACC |
| Core | GATAAGCGTAAGGTTATCTAAAATATCGGATTTAGAG |
| Core | ACACAAGAGTGGTTTAATATCATCGCCTG |
| Core | ATTTTATCCTGAAGCCTTAAAGTTTTTATAACGGGTA |
| Core | GGGTGAACGGTAATCGTAAACGTCGGATAATAATTCCCA |
| Core | GTAAACTATCGCAGTAATAAGCGGGAGCTAAACAG |
| Core | TCTCTGAATTTACAAACAAATGCCGCCACCAA |
| Core | AGACGATTCAGCACCGTAATCAGTTCACCGACTCAACCGAGTG |
| Core | CCCTCATACATGTACCAAAAAGGCTCCAAAAGGCGC |
| Core | TGTAGGTAAAGAAACGCAAGGGCAAAGATGAAAAGG |
| Core | CCAGCAAAAGAATAGCCCGAGTTGAGTGTTGTTCCA |
| Core | ATACTTGAGATAATCTTGATGAAAGAGGACA |
| Core | CGAACCAGAATTCTACCATATAACATTTTTAGAACCCTCA |
| Core | CGGAATTGTTTGGATTATACATTTGAATACCAAGTAGG |
| Core | AACGTCAAAAATGCATTACCGCCCGACTTAACGCGCC |
| Core | TGAATGGCGGCAAATCAACAGTTGTCGACAACATCA |
| Core | TGTTTATCGGCATTTTCGCTCAACTGTGATAAATAAGGCGTAAT |
| Core | TTAATCATTGTGATAGAATCATAGGCTGGCTGACCTTCATTTCA |
| Core | TTCCTGGATAGAGAGCAACTTTAGGAATACC |
| Core | GAATCTTACCAGCCTTTACACCCTGAACA |
| Core | AATAAAATAGCAACGCTAAATAGCTATCTTA |
| Core | GAAAAACAGAAGCAAATGAAAAATCACCAGAAGGAAG |
| Core | TGGGGAGTAGATGAGAAGCCTTTATTTCTTCAAAAGGAGA |
| Core | TTTTGGGTTAGAACCTACCAAAGAAAACATTCATTTAGA |
| Core | GAAACGTCACCAACAGCAAAATTTACCAGCAC |
| Core | CACCAGCAATATTACTTTACATTGATTTTAGACAGGAAC |
| Core | TTAATTGAATGTAGAAATAAGTCCAACGCGAGGCGTT |
| Core | ATCATAACCCTCGCATCAAAAAGAATGACTATAATGC |
| Core | TAATAAAATCATTACAAAATCGCGCAGAGTAGATTTTAATGGAAGCG |
| Core | GCTGTAACCAATAGGAACGAAACGTTAGAATCGATAGC |

| | |
|---|---|
| Core | CCCACGCGTTGAAAAAATAGAAAAGCATTCCACAGACAG |
| Core | TACCAGAAGGAAAAAGCCCTTTTAACTGAACAGAGAG |
| Core | TGTAGCTCCTGCGAACGCGCGAGCATTAGCAAAATTAAGCTAAT |
| Core | TACCAAGCGCGAACATCACAGGGT |
| Core | CGGGTTGCAACAGTGCTGATTATCGCCGTCAATTTACATC |
| Core | TCTGAAACATGAACCCTCAGAAAATCCTCCCG |
| Core | ACCATTAGGAGCTTAACTTTAATTGAAGCAAAGCGGATTGTTTACCAGGG |
| Core | TTGGTAGTAAAATGTTTAGAAGAAAACGAGAT |
| Core | TATCAGTTAGTTGGTTTGCCCCAGCAGGCGAA |
| Core | TCCCAATTAACATGTTGCAAACTCAGGAAGCCCGAA |
| Core | CAGTGCCTTGAGTAACGAACCACCCCGCCAGCTTGCCTTT |
| Core | CCGGTTGAGAAAAGCCCCAAAAACTAGCCAGCATTAAATGCTCAGGAA |
| Core | TCACATTACGCGCGGCTTCACCGTGGTTCCGAAATCGGC |
| Core | CTAATGAGCAAGGCGATTAAGTTGCTTCTGGTGTATCGGCTGA |
| Core | ACTGTTGGGAAGGGCGAAGAAGAACGTTGGTGTAGA |
| Core | GGACTCCATTTGATGGCCTGGCCCTGAGAGAATGG |
| Core | GATAAGAAACGTGCCAGTTCAATAATAAGAGCAAGAAAGAACTACAT |
| Core | GCACAGACATGGATTACGCCAGCCGCAAATTAACCGTTGT |
| Core | CTAGCTGATAAATAATAAAGCTTTTGCGGTTAGTTTG |
| Core | TTATCCCAATCCAAATATAGAAGGATTCTAAGTGAACAAG |
| Core | TTTAGTAAGTACCGCCCAGACGAAGTTGCTATTTATTCA |
| Core | GAGGCCGTTCTTTGGGTAATATCCAGAACAGAA |
| Core | AAATAGCGTTTTGCAAAAGAAGTTGGTCTTTATATAGTCAGCTCCTTT |
| Core | ATCCTTGAAAACATTTGAAATTCAAATATATGTAATT |
| Core | ATTAGAAATAAAGAAATTGCGGCGAATTAAAATTAAAGA |
| Core | GCGCGTTTTCATCGGCAAATATTGTTCATTAATAAGACTC |
| Core | GATGCTTTGAGACCCCCTTGATACCGATAGTTGAGC |
| Core | CCCGGAATTGGTAATAAGTTTTAA |
| Core | CCACCCTCTTCCAGACGTTAGGCT |
| Core | CATCGGAACAAACGGCGGAAACAAGAATATTTTATG |
| Core | CTGCACACGACGCCTTGCTATTAGTAATAACATCA |
| Core | TCACCAGTACGCCTGTGGAACAACTAAAGGAAATAATAAT |
| Core | TAAGCAACAGGTCAGGATTACATT |
| Core | TCCTCGCTCAATCGTCTGAAAATATTTT |
| Core | AGTAGTTAAAATTCGCATTAAATTTAGACAAATTGC |
| Core | AAAAATACTGAACCTCAAATATCAGTAACATTTCGT |
| Core | CGACTTGCGGGAGACTTTTTCATGAGGACCGAACTGCGAA |
| Core | GCCGGGTGAGAACAAATATTTAAATTGTCCATCAAATCTCCGTGTGC |
| Core | GAATTTTCTGTATGGGGCGAAAGATAAACAGCAGCGATTA |
| Core | CTGAGAGTCTGGAGCATTGACCGTTCATTTTTCGCA |
| Core | AAGCCATCTTTATCGATAGGGCCTTGATATTCACCGTT |
| Core | GTTTGGAATTATAAATTGAGACGGGCAACAGCTATCCGCT |
| Core | ACACAAGAGTGGTTTAATATCATCGCCTG |

| | |
|---|---|
| Core | ATTTTATCCTGAAGCCTTAAAGTTTTTATAACGGGTA |
| Core | GGGTGAACGGTAATCGTAAACGTCGGATAATAATTCCCA |
| Core | GTAAACTATCGCAGTAATAAGCGGGAGCTAAACAG |
| Core | TCTCTGAATTTACAAACAAATGCCGCCACCAA |
| Core | AGACGATTCAGCACCGTAATCAGTTCACCGACTCAACCGAGTG |
| Core | CCCTCATACATGTACCAAAAAGGCTCCAAAAGGCGC |
| Core | TGTAGGTAAAGAAACGCAAGGGCAAAGATGAAAAGG |
| Core | CCAGCAAAAGAATAGCCCGAGTTGAGTGTTGTTCCA |
| Core | ATACTTGAGATAATCTTGATGAAAGAGGACA |
| Core | CGAACCAGAATTCTACCATATAACATTTTTAGAACCCTCA |
| Core | CGGAATTGTTTGGATTATACATTTGAATACCAAGTAGG |
| Core | AACGTCAAAAATGCATTACCGCCCGACTTAACGCGCC |
| Core | TGAATGGCGGCAAATCAACAGTTGTCGACAACATCA |
| Core | TGTTTATCGGCATTTTCGCTCAACTGTGATAAATAAGGCGTAAT |
| Core | TTAATCATTGTGATAGAATCATAGGCTGGCTGACCTTCATTTCA |
| Core | TTCCTGGATAGAGAGCAACTTTAGGAATACC |
| Core | GAATCTTACCAGCCTTTACACCCTGAACA |
| Core | AATAAAATAGCAACGCTAAATAGCTATCTTA |
| Core | GAAAAACAGAAGCAAATGAAAAATCACCAGAAGGAAG |
| Core | TGGGGAGTAGATGAGAAGCCTTTATTTCTTCAAAAGGAGA |
| Core | TTTTGGGTTAGAACCTACCAAAGAAAACATTCATTTAGA |
| Core | GAAACGTCACCAACAGCAAAATTTACCAGCAC |
| Core | CACCAGCAATATTACTTTACATTGATTTTAGACAGGAAC |
| Core | TTAATTGAATGTAGAAATAAGTCCAACGCGAGGCGTT |
| Core | ATCATAACCCTCGCATCAAAAAGAATGACTATAATGC |
| Core | TAATAAAATCATTACAAAATCGCGCAGAGTAGATTTTAATGGAAGCG |
| Core | GCTGTAACCAATAGGAACGAAACGTTAGAATCGATAGC |
| Core | CCCACGCGTTGAAAAAATAGAAAAGCATTCCACAGACAG |
| Core | TACCAGAAGGAAAAAGCCCTTTTAACTGAACAGAGAG |
| Core | TGTAGCTCCTGCGAACGCGCGAGCATTAGCAAAATTAAGCTAAT |
| Core | TACCAAGCGCGAACATCACAGGGT |
| Core | CGGGTTGCAACAGTGCTGATTATCGCCGTCAATTTACATC |
| Core | TCTGAAACATGAACCCTCAGAAAATCCTCCCG |
| Core | ACCATTAGGAGCTTAACTTTAATTGAAGCAAAGCGGATTGTTTACCAGGG |
| Core | TTGGTAGTAAAATGTTTAGAAGAAAACGAGAT |
| Core | TATCAGTTAGTTGGTTTGCCCCAGCAGGCGAA |
| Core | TCCCAATTAACATGTTGCAAACTCAGGAAGCCCGAA |
| Core | CAGTGCCTTGAGTAACGAACCACCCCGCCAGCTTGCCTTT |
| Core | CCGGTTGAGAAAAGCCCCAAAAACTAGCCAGCATTAAATGCTCAGGAA |
| Core | TCACATTACGCGCGGCTTCACCGTGGTTCCGAAATCGGC |
| Core | CTAATGAGCAAGGCGATTAAGTTGCTTCTGGTGTATCGGCTGA |
| Core | ACTGTTGGGAAGGGCGAAGAAGAACGTTGGTGTAGA |
| Core | GGACTCCATTTGATGGCCTGGCCCTGAGAGAATGG |

| | |
|---|---|
| Core | GATAAGAAACGTGCCAGTTCAATAATAAGAGCAAGAAAGAACTACAT |
| Core | GCACAGACATGGATTACGCCAGCCGCAAATTAACCGTTGT |
| Core | CTAGCTGATAAATAATAAAGCTTTTGCGGTTAGTTTG |
| Core | TTATCCCAATCCAAATATAGAAGGATTCTAAGTGAACAAG |
| Core | TTTAGTAAGTACCGCCCAGACGAAGTTGCTATTTATTCA |
| Core | GAGGCCGTTCTTTGGGTAATATCCAGAACAGAA |
| Core | AAATAGCGTTTTGCAAAAGAAGTTGGTCTTTATATAGTCAGCTCCTTT |
| Core | ATCCTTGAAAACATTTGAAATTCAAATATATGTAATT |
| Core | ATTAGAAATAAAGAAATTGCGGCGAATTAAAATTAAAGA |
| Core | GCGCGTTTTCATCGGCAAATATTGTTCATTAATAAGACTC |
| Core | GATGCTTTGAGACCCCCTTGATACCGATAGTTGAGC |
| Core | CCCGGAATTGGTAATAAGTTTTAA |
| Core | CCACCCTCTTCCAGACGTTAGGCT |
| Core | CATCGGAACAAACGGCGGAAACAAGAATATTTTATG |
| Core | CTGCACACGACGCCTTGCTATTAGTAATAACATCA |
| Core | TCACCAGTACGCCTGTGGAACAACTAAAGGAAATAATAAT |
| Core | TAAGCAACAGGTCAGGATTACATT |
| Core | TCCTCGCTCAATCGTCTGAAAATATTTT |
| Core | AGTAGTTAAAATTCGCATTAAATTTAGACAAATTGC |
| Core | AAAAATACTGAACCTCAAATATCAGTAACATTTCGT |
| Core | CGACTTGCGGGAGACTTTTTCATGAGGACCGAACTGCGAA |
| Core | GCCGGGTGAGAACAAATATTTAAATTGTCCATCAAATCTCCGTGTGC |
| Core | GAATTTTCTGTATGGGGCGAAAGATAAACAGCAGCGATTA |
| Core | CTGAGAGTCTGGAGCATTGACCGTTCATTTTTCGCA |
| Core | AAGCCATCTTTATCGATAGGGCCTTGATATTCACCGTT |
| Core | GTTTGGAATTATAAATTGAGACGGCAACAGCTATCCGCT |
| Top-ext A | CTTTACAAACAATAAAGGAATCACCTTGCCGAACGAACTTTCTCTCTCTCTC |
| Top-ext B | GGCAAAGCGCCATAAAGGGGGCCAAGCTTTCCTGTGTTTTCTCTCTCTCTC |
| Top-ext C | ACCTGCTCCATGTAACGAAAGTTAAACGGCTTGCAGGTTTCTCTCTCTCTC |
| Top-ext D | TGATAGGTGAATTATCACCGAGCGACAGTCATAGCCCTCAGAGCTTTCTCTCTCTCTC |
| Top-ext I | TTCGGAAGTTTCATTCTAATAGTAGTAGCCCTGTTTCTCTCTCTCTC |
| Bott-ext A | GAACAAAGAAACCTAAAGCATTGAGGAAGAATACGTGTTTAGGAGGAGGAGGAACAG CCC |
| Bott-ext B | CAGTTTGAGGGGAGGCCAGTGATGTGCTGTGAGCTAACTTTAGGAGGAGGAGGAACAG CCC |
| Bott-ext C | AATCATAAGGGAAAGTTTCCAAGGCAAAACAACCATCGTTTAGGAGGAGGAGGAACA GCCC |
| Bott-ext D | AAAGTTGAGGGAGGGAAGGTATTTTCGGAATCAAGTATTGACAGGTTTAGGAGGAGGA GGAACAGCCC |
| Bott-ext I | AGGTCTATATGTGAGTACTAGAAATATAACTATAAAGTAATTTAGGAGGAGGAGGAAC AGCCC |

**Table S6. Parameters derived from the Gaussian mixture model decomposition.**

The segments were measured across the different stamps and the distributions were decomposed using a two-component Gaussian mixture model. The number of input observations (n) yielded the weights (w1 and w2) as well as means (mean1, mean2) and variances (var1, var2) for each Gaussian component respectively.

| *Sample* | Section | n | w1 | w2 | mean1 | mean2 | var1 | var2 |
|---|---|---|---|---|---|---|---|---|
| *1-step Biotin* | AB | 4562 | 0.69 | 0.31 | 29.79 | 40.93 | 5.13 | 15.89 |
| | AD | 4532 | 0.90 | 0.10 | 44.81 | 30.82 | 11.74 | 7.92 |
| | BI | 4725 | 0.66 | 0.34 | 22.69 | 30.64 | 9.88 | 19.56 |
| | CB | 4727 | 0.82 | 0.18 | 41.89 | 37.62 | 5.23 | 44.77 |
| | DC | 4700 | 0.10 | 0.90 | 30.93 | 42.21 | 6.52 | 8.00 |
| | ID | 4697 | 0.64 | 0.36 | 33.65 | 26.18 | 12.44 | 13.23 |
| *3x Biotin* | AB | 3207 | 0.27 | 0.73 | 41.28 | 29.77 | 10.26 | 3.97 |
| | AD | 3193 | 0.10 | 0.90 | 30.47 | 44.13 | 5.07 | 8.40 |
| | BI | 3300 | 0.30 | 0.70 | 29.87 | 23.01 | 19.11 | 7.07 |
| | CB | 3308 | 0.16 | 0.84 | 36.88 | 41.68 | 43.97 | 4.69 |
| | DC | 3293 | 0.91 | 0.09 | 41.91 | 30.75 | 7.41 | 5.84 |
| | ID | 3285 | 0.37 | 0.63 | 26.89 | 32.67 | 11.78 | 8.45 |
| *3x Biotin-7T* | AB | 790 | 0.23 | 0.77 | 39.62 | 29.95 | 25.94 | 9.02 |
| | AD | 782 | 0.28 | 0.72 | 41.94 | 44.92 | 79.58 | 8.20 |
| | BI | 826 | 0.71 | 0.29 | 22.68 | 29.55 | 13.17 | 19.27 |
| | CB | 826 | 0.27 | 0.73 | 38.34 | 41.04 | 38.86 | 6.33 |
| | DC | 818 | 0.75 | 0.25 | 41.27 | 39.39 | 6.23 | 46.70 |
| | ID | 818 | 0.57 | 0.43 | 34.38 | 27.99 | 19.38 | 19.60 |
| *2-step Biotin* | AB | 2731 | 0.79 | 0.21 | 29.28 | 40.26 | 6.10 | 15.37 |
| | AD | 2725 | 0.89 | 0.11 | 44.24 | 31.01 | 9.91 | 8.09 |
| | BI | 2959 | 0.77 | 0.23 | 20.37 | 30.75 | 9.12 | 26.05 |
| | CB | 2999 | 0.23 | 0.77 | 37.00 | 40.86 | 33.99 | 5.42 |
| | DC | 2991 | 0.18 | 0.82 | 39.82 | 42.03 | 50.17 | 3.85 |
| | ID | 2952 | 0.23 | 0.77 | 24.85 | 35.43 | 22.05 | 12.57 |
